# Dynamic functional connectivity encodes generalizable representations of emotional arousal across individuals and situational contexts

**DOI:** 10.1101/2023.11.14.566767

**Authors:** Jin Ke, Hayoung Song, Zihan Bai, Monica D. Rosenberg, Yuan Chang Leong

## Abstract

Human affective experience varies along the dimensions of valence (positivity or negativity) and arousal (high or low activation). It remains unclear how these dimensions are encoded in the brain and if the representations are shared across different individuals and diverse situational contexts. Here we utilized two publicly available functional MRI datasets of participants watching movies to build predictive models of moment-to-moment valence and arousal from dynamic functional brain connectivity. We tested the models both within and across datasets and identified a generalizable arousal representation characterized by the interaction between multiple large-scale functional networks. The arousal representation generalized to two additional movie-watching datasets. Predictions based on multivariate patterns of activation underperformed connectome-based predictions and did not generalize. In contrast, we found no evidence of a generalizable valence representation. Taken together, our findings reveal a generalizable representation of arousal encoded in patterns of dynamic functional connectivity, revealing an underlying similarity in how arousal is encoded across individuals and situational contexts.

## Introduction

Human experience is characterized by a continual ebb and flow of emotions, shaping our attention (*1*, *2*) and memory (*3*, *4*), as well as guiding how we make decisions and interact with others (*5–7*). These affective experiences, though diverse and complex, are often thought to be organized along a small number of principal dimensions (*8–10*). Among these, valence and arousal are two dimensions that are central to many contemporary theories of emotion (*11–14*). Valence refers to the positivity or negativity of an emotional state, while arousal refers to its intensity or activation level. Emotional states can be placed in a two-dimensional space based on their level of valence and arousal—for example, both “excitement” and “anger” are high in arousal but occupy opposite ends on the valence dimension (*15*). Thus, this framework provides a useful model for illustrating how seemingly distinct emotional experiences share underlying similarities and relate to one another in a structured space. A recent study found that individuals’ positions within this space in a given situation predicts their behavior during social interactions (*16*), suggesting that these dimensions not only describe emotional states but also serve as reliable indicators for predicting behavioral responses.

The concept of valence and arousal as core components of affect has significantly influenced research and theories in emotion (*17–19*). It has been proposed that valence and arousal are not inherent attributes of emotional states themselves, but are the outcomes of individual appraisals or interpretations of an event, shaped by past experiences, contextual information, and sensory input (*14*, *20–22*). This raises a crucial question about the nature of how emotions are represented in the brain: Are the neural representations of valence and arousal consistent across different individuals and situations? One possibility is that these representations are specific to each situation and each individual, meaning that the brain encodes experiences with similar levels of valence or arousal differently depending on the context or person. Alternatively, there could be generalizable neural patterns that consistently encode valence or arousal across various contexts and individuals. For example, the brain might encode arousal from an exciting movie scene and from an anger-inducing interpersonal interaction similarly. Such generalizability in neural representations might explain why people sometimes mistakenly attribute their arousal to incorrect causes (*23–25*). Determining whether valence and arousal are encoded in a generalizable manner contributes to a deeper understanding on the neural basis of affective experiences.

Prior research has identified several brain regions involved in encoding valence and arousal, including the orbitofrontal cortex (OFC), amygdala, ventral striatum, and insula (*26–31*). A seminal study by Chikazoe and colleagues (*32*) identified a multivariate representation of valence in OFC activity that generalizes across both visual and gustatory stimuli, as well as across different individuals, suggesting a common neural code for subjective valence. However, it remains less clear whether a similarly generalizable neural code exists for arousal. Furthermore, previous studies have often relied on brief, decontextualized stimuli, such as isolated visual images or words, to investigate these affective dimensions (*26–32*). Such stimuli may not capture the diversity and complexity of real-world emotional experiences, raising questions about whether the neural representations of valence and arousal identified under these controlled conditions would generalize across varied and dynamic real-life situations, and across different individuals encountering these contexts.

Existing work on the neural basis of valence and arousal have primarily focused on overall activity or activity patterns within specific brain regions (*26–33*). This approach, while foundational, does not consider the role of dynamic interactions between large-scale functional brain networks. Affective experience involves the integration of sensory inputs, past experiences and current goals, which requires the coordination between networks with different functions (*34*). Indeed, there is growing recognition that whole-brain interactions play an important role in facilitating diverse cognitive processes (*35–37*), with recent studies demonstrating that whole-brain functional connectivity patterns can predict both intra-individual fluctuations and inter-individual differences in attention (*38, 39*), memory (*40, 41*) and comprehension of complex narratives (*42*). The relationship between whole-brain functional connectivity patterns and affective experience, however, remains largely unexplored.

The goal of the present study is to test whether generalizable neural representations of valence and arousal are encoded in the interactions between large-scale functional brain networks. Our work has three key features. First, we trained predictive models (*43*) to predict the valence and arousal of viewed stimuli from whole-brain dynamic functional connectivity patterns. We used the dynamic connectome-based predictive modeling (CPM) approach developed by Song et al. (*39*, *42*) to predict moment-to-moment ratings of valence and arousal from whole-brain functional connectivity dynamics. CPMs are so named because they leverage the connectome (i.e., the comprehensive map of functional connections in the brain) to make predictions about inter- or intra-individual differences in behavior or mental states. Second, rather than presenting participants with static images or short video clips designed to elicit a specific affective state, we examined continuous affective responses to full-length TV episodes. These episodes have comprehensive narrative arcs that elicited a wide range of affective responses (*39*, *44*, *45*) and allowed us to capture the moment-to-moment affective responses across a variety of situations. Third, we assessed the out-of-sample generalizability (*46*) of our findings by testing our models on new stimuli and participants that were not used to train the model. This step is crucial to ascertain whether our models were fitting to the idiosyncrasies of specific contexts and a group of individuals, or if they genuinely reflect generalizable neural representations of valence and arousal.

We utilized two publicly available fMRI datasets of participants watching TV episodes (*N* = 16 (*44*) and *N* = 35 (*45*); Fig. 1), and collected continuous valence and arousal ratings of the two episodes from a separate group of participants (*N* = 120, 30 in each condition). We built CPMs to predict valence and arousal fluctuations from dynamic functional connectivity patterns. We assessed both the within-dataset accuracy (i.e., how well the CPM predicted valence or arousal in the dataset it was trained on) as well as across-dataset accuracy (i.e., how well a CPM predicted valence or arousal in the dataset it was not trained on). To further validate the robustness and generalizability of our results, we tested the trained CPMs on two additional fMRI datasets of different participants watching different movies. Additionally, we repeated our analyses with univariate and multivariate methods to compare the predictive accuracy of the CPM against more traditional analytical approaches. Altogether, our approach seeks to identify generalizable neural representations of moment-to-moment valence and arousal based on the interaction between brain areas, and shed new light on how affective experiences are encoded in the brain.

**Fig. 1.**
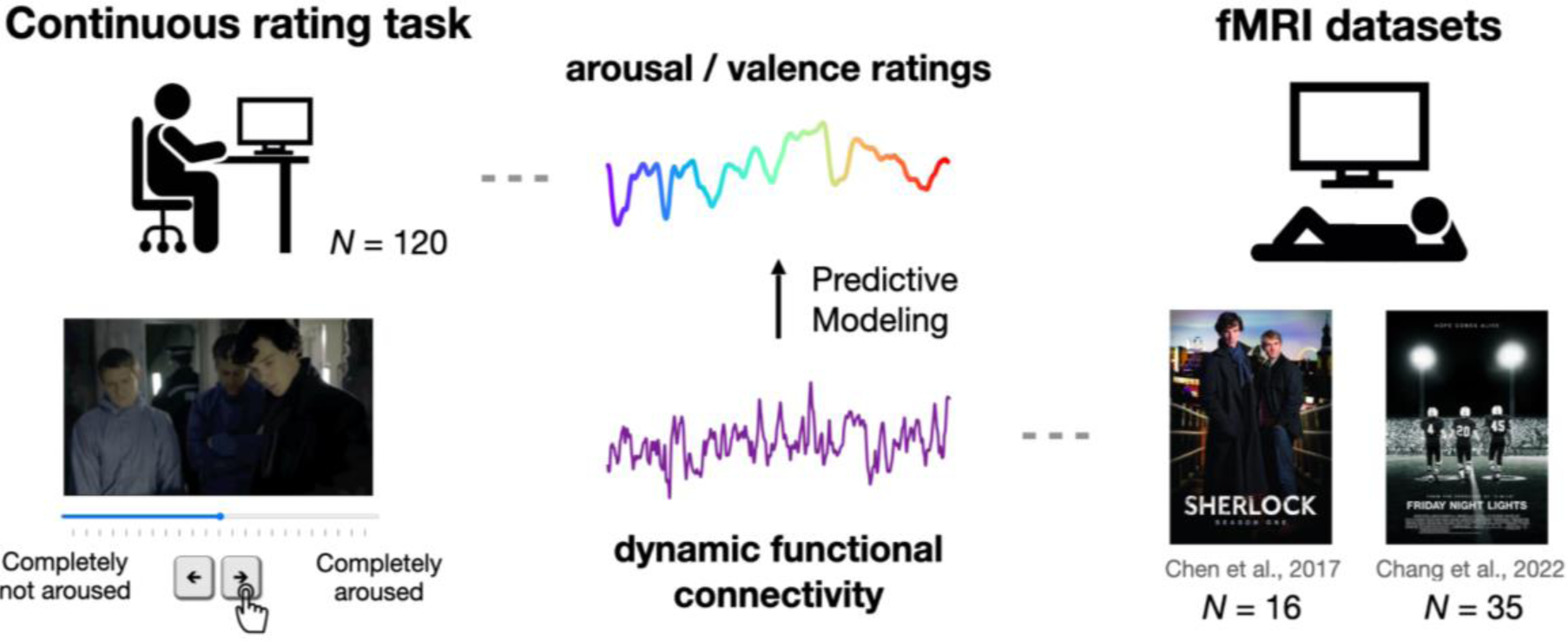
Study schematic procedure. We utilized two publicly available fMRI datasets of participants watching *Sherlock* (*N* = 16) and *Friday Night Lights* (*N* = 35). Additionally, we collected behavioral ratings of the two episodes from a separate group of participants (*N* = 120). Each participant is presented with one movie clip and is instructed to continuously rate how the clip is making them feel in terms of either valence (i.e., positive to negative) or arousal (i.e., not aroused to aroused). We built within and across dataset CPMs to predict moment-to-moment affect ratings from dynamic whole-brain functional connectivity patterns.

## Results

### Affective experience is synchronized across individuals during movie watching

One-hundred and twenty participants performed an affect rating task while watching one of two TV episodes. One was a 48-min long episode from BBC’s *Sherlock* (a British mystery crime drama series) while the other was a 45-min long episode from NBC’s *Friday Night Lights* (an American sports drama series). Each participant provided continuous ratings of either valence (i.e., positive to negative) or arousal (i.e., not aroused to aroused) while watching one of the videos (**Fig. 1**). As we were primarily interested in subjective feelings during the movie, we instructed participants to indicate their affective experience (e.g., how positive or negative they were feeling), rather than the perceived emotionality of the movies (e.g., whether they thought the current scene was positive or negative). Previous work found that providing continuous affect ratings did not alter emotional or neural responses to emotion-eliciting films, indicating that this approach can capture participants’ affective experiences without disrupting their natural responses to the content (*47*). We will refer to the pairing of an affective dimension and a movie as an “experimental condition” (e.g., *Sherlock*-valence). Thus, we have four experimental conditions with a sample size of 30 participants each. The group-averaged time courses were treated as a proxy of the affective experience time-locked to the movie (**Fig. 2A**).

**Fig. 2.**
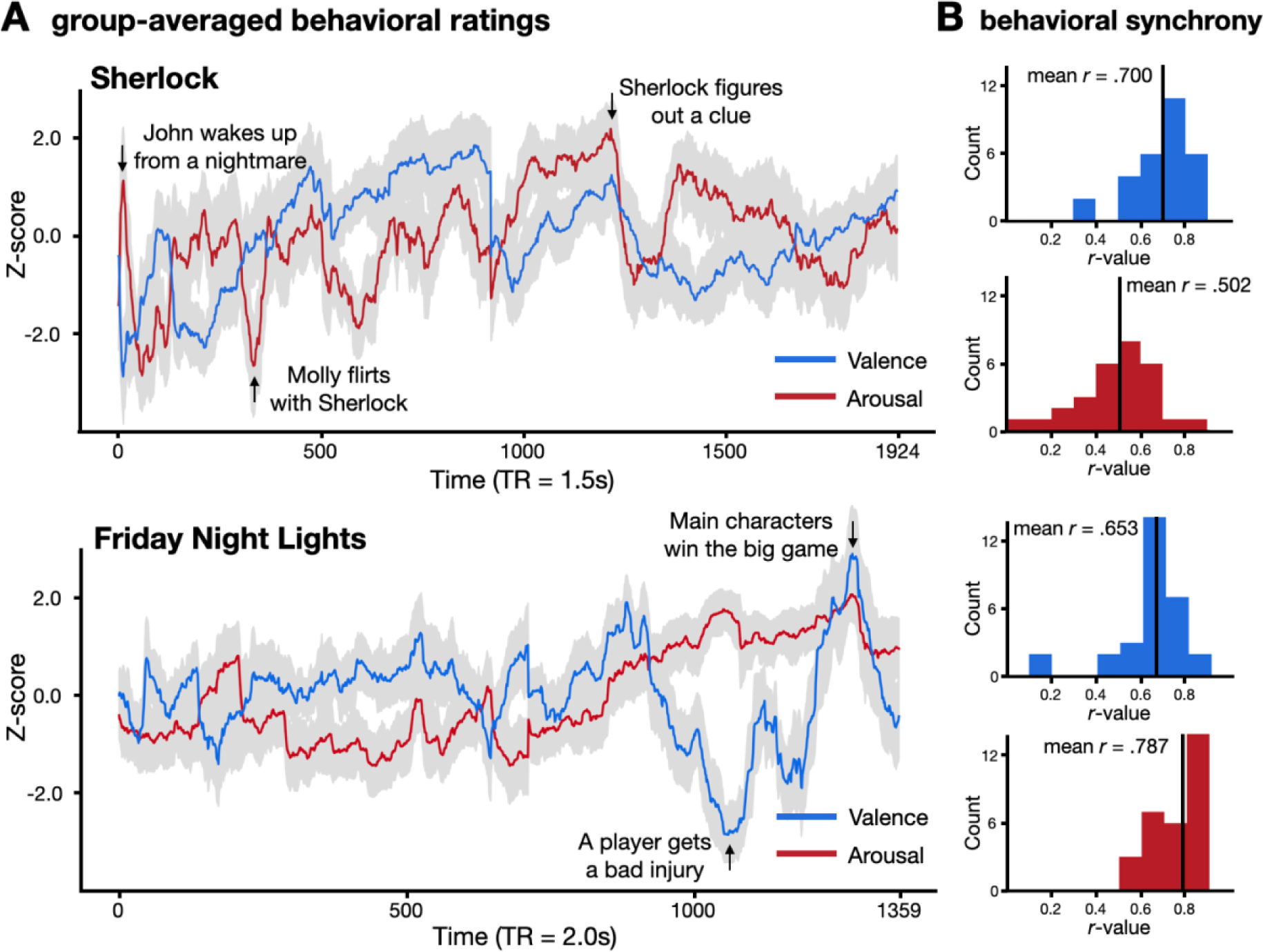
Affective experience is synchronized across participants in all four experimental conditions. Each condition includes 30 participants. **A.** Participants’ subjective affective experience fluctuates over time during naturalistic movie watching. The red and blue solid lines indicate average arousal and valence time courses respectively. The gray areas indicate 95% confidence interval (CI). **B.** Histograms of the similarity of each individual’s subjective ratings to the group-averaged rating with the individual left out. Higher mean *r* values indicate stronger affective synchrony.

We first examined whether subjective ratings of valence and arousal were synchronized across individuals during movie watching. We computed the Pearson correlation between each individual’s rating and the group-averaged rating with the individual left out. The distribution of intersubject correlations was significantly positive in all conditions (**Fig. 2B**; *Sherlock-*valence: mean *r* = .700, *s.d.* = .197, *p* < .001; *Sherlock-*arousal: mean *r* = .502, *s.d.* = .196, *p* < .001; *Friday Night Lights*-valence: mean *r* = .653, *s.d.* = .167, *p* < .001; *Friday Night Lights-*arousal: mean *r* = .787, *s.d.* = .109, *p* < .001). Statistical significance was assessed with a nonparametric permutation test (*48*). These results suggest that affective experience, across both valence and arousal dimensions, was synchronized across individuals when they watched both TV episodes.

Qualitatively, the average valence and arousal time courses reflected expected fluctuations in affective experience during the episodes (**Fig. 2A**). For example, while watching *Sherlock*, participants’ ratings indicated high negative valence and high arousal during the scene where John Watson has a nightmare of his time in the war, and high positive valence but low arousal during a scene where Sherlock was clueless to the lab assistant, Molly, flirting with him. Similarly, while watching *Friday Night Lights*, participants’ ratings indicated high negative valence and high arousal when a star player is severely injured in a football game, and high positive valence and high arousal when the main character’s team wins the game. The average valence and arousal time courses were not significantly correlated in both *Sherlock* (*r* = .094, *p* = .359) and *Friday Night Lights* (*r* = -.224, *p* = .187). The average time courses of arousal and the absolute value of valence were significantly correlated in *Friday Night Lights* (*r* = .706, *p* < .001) but not *Sherlock* (*r* = .101, *p* = .303).

### Dynamic functional connectivity encodes arousal within and across datasets

We first asked whether dynamic functional connectivity encoded fluctuations in arousal. To that end, we trained CPMs to predict arousal rating time courses from time-resolved dynamic functional connectivity (FC) patterns (**Fig. 1**). We parcellated the whole brain into 122 ROIs, following the Yeo atlas for cortical regions (*49*) (114 ROIs) and the Brainnetome atlas for subcortical regions (*50*) (8 ROIs: bilateral amygdala, hippocampus, basal ganglia, and thalamus). The dynamic FC patterns were extracted from the 122-ROI-based BOLD time courses using a tapered sliding window approach, where the Fisher’s z-transformed Pearson’s correlation between the BOLD time courses of every pair of ROIs was computed within each tapered sliding window (*39*) (window size - *Sherlock*: 30 TRs = 45s; *Friday Night Lights*: 23 TRs = 46s; see Methods).

Separate models were trained on two openly available fMRI datasets where participants watched *Sherlock* (*44*) (*N* = 16) or *Friday Night Lights* (*45*) (*N* = 35). We assessed model performance in predicting group-mean arousal ratings within each dataset as well as between datasets. *Within-dataset performance* was computed using a leave-one-subject-out cross-validation approach, training the model on the neural data of all but one participant in a dataset, and applying the trained model to data from the held-out participant to predict the average arousal time course. In every round of cross-validation we selected FCs that significantly correlated with arousal in the training set as features (one-sample Student’s *t* test, *p* < .01). Model accuracy was computed as the average correlation between the model-predicted arousal time course and the empirical arousal time course across cross-validation folds. Statistical significance was assessed by comparing model accuracy against a null distribution of 1000 permutations generated by training and testing the model on phase-randomized behavioral ratings. We note that the null distributions for within-dataset performance were positively skewed, as each iteration of the permutation test involved training and testing on the same phase-randomized behavioral rating. This meant that the null model could learn associations between FC connections and arbitrary stimulus features that correlated with the randomized engagement time course. Thus, the null distribution provided a conservative test of model performance against a baseline measure of chance performance due to non-specific associations in the data.

Within-dataset accuracy was significantly above chance in predicting arousal ratings for both *Sherlock* (mean *r* = .575, *p* = .034; MSE = .680, *p* = .034; R^2^ = .320, *p* = .035) and *Friday Night Lights* (mean *r* = .734, *p* = .002; MSE = .491, *p* < .001; R^2^ = .509, *p* = .002; **Fig. 3A**, left), suggesting that dynamic whole-brain FC patterns encoded moment-to-moment experience of arousal during both movies. However, an alternative possibility was that the CPM was learning features specific to the particular movie it was trained on rather than arousal per se. For example, if Sherlock Holmes tended to appear in scenes that were highly arousing, the model could learn to associate the neural representation of Sherlock Holmes with higher arousal ratings. In a different movie, where Sherlock Holmes was not present, or no longer associated with arousing scenes, the model would fail to predict arousal. Thus, an important test of whether the model encodes arousal would require testing the model in a novel context.

**Fig. 3.**
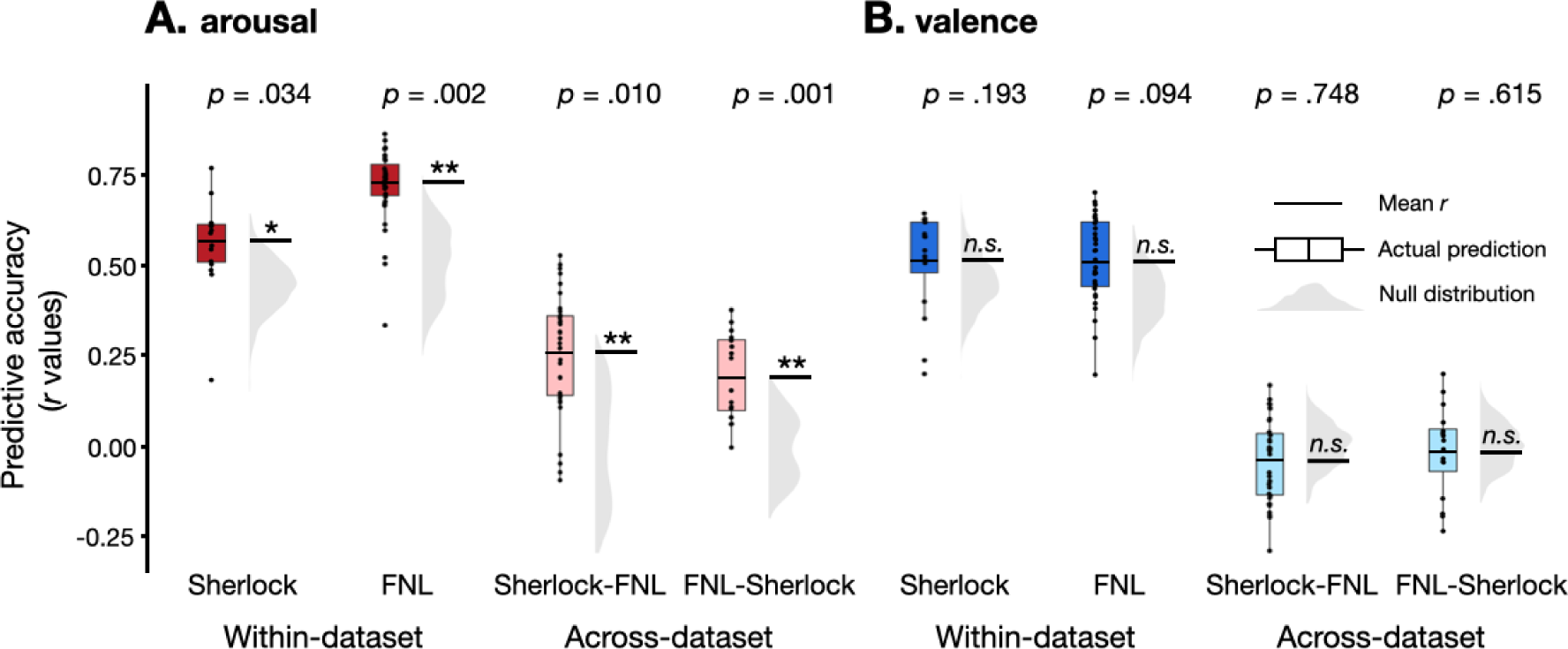
Dynamic functional connectivity predicts fluctuations in arousal but not valence. CPM performance in predicting arousal (**A**) and valence (**B**) for within-dataset (the left panel) and between-dataset (the right panel). The y-axis represents the predictive accuracy, as measured by Pearson’s correlation between the model predicted time course and the observed group-average time course. Each datapoint in the box plot represents the predictive accuracy of each round of cross-validation. The black horizontal lines show the mean *r* value computed from the average of the Fisher-*z* transformed individual-subject *r* values. The gray half-violin plots show the null distribution of 1000 permutations, generated by phase-randomizing the observed group-average time course before training and testing the models. \**p* < 0.05, \*\**p* < 0.01, n.s.: p > 0.05, as assessed by comparing the empirical mean predictive accuracy against the null distribution.

With that goal in mind, we assessed the *across-dataset performance* to test whether the models generalized across datasets. For each dataset, we first identified the set of FC features that were selected in every round of the cross-validation, which we term the arousal network for that dataset. To test model generalizability, we trained a CPM in one dataset on its arousal network to predict the group-average arousal time course and applied the trained model to the other dataset with a different group of participants watching a different movie. Here, the null distributions were centered around zero, because the two datasets featured different stimuli. Thus, any random associations learned by the model from the training dataset are unlikely to apply to the testing dataset due to the lack of overlap in the stimuli. The across-dataset predictions of arousal (**Fig. 3A**, right) were significantly above chance both for a model trained on *Sherlock* and tested on *Friday Night Lights* (mean *r* = .270, *p* = .010; MSE = .954, *p* = .009; R^2^ = .046, *p* = .008), and a model trained on *Friday Night Lights* and tested on *Sherlock* (mean *r* = .198, *p* = .001; MSE = .998, *p* = .001; R^2^ = .002, *p* = .001).

To assess whether prediction results were driven by low-level audio-visual features, we reran the prediction analyses after regressing out ten low-level features (hue, saturation, pixel intensity, motion energy, presence of a face, whether the scene was indoor or outdoor, presence of written text, amplitude, pitch, presence of music) from each participant’s BOLD signal time courses. These features were selected following the procedures prior studies examining the neural correlates of high-level cognitive processes using dynamic audio-visual stimuli (*51*, *52*). Consistent with our previous analyses, we found above-chance predictions when the models were tested within each dataset (*Sherlock*: mean *r* = .558, *p* = .024; MSE = .696, *p* = .025; R^2^ = .303, *p* = .026 ; *Friday Night Lights*: mean *r* = .732, *p* = .003; MSE = .491, *p* = .002; R^2^ = .509, *p* = .003) and across datasets (train on *Sherlock* test on *Friday Night Lights*: mean *r* = .267, *p* = .012; MSE = .959, *p* = .011 ; R^2^ = .040, *p* = .012; train on *Friday Night Lights* test on *Sherlock*: mean *r* = .220, *p* = .020; MSE = .981, *p* < .001; R^2^ = .018, *p* < .001), suggesting that prediction accuracy was not confounded by these low-level features.

Additionally, to examine whether motion influenced the CPM predictions on arousal, we re-ran the analyses after regressing out each individual’s frame-to-frame displacement from their dynamic functional connectivity time courses. Consistent with our previous results, the model significantly predicted arousal in both *Sherlock* (mean *r* = .571, *p* = .028.; MSE = .684, *p* = .030; R^2^ = .316, *p* = .032) and *Friday Night Lights* (mean *r* = .641, *p* = .004; MSE = .624, *p* = .015; R^2^ = .376, *p* = .018). The across-dataset predictions were similarly significantly above chance both when trained on *Sherlock* tested on *Friday Night Lights* (mean *r* = .238, *p* = .015; MSE = .980, *p* = .025.; R^2^ = .020, *p* = .026) and trained on *Friday Night Lights* tested on *Sherlock* (mean *r* = .151, *p* = .034; MSE = 1.014, *p* = .036; R^2^ = -.015, *p* = .037). These results suggest that our results were unlikely to be driven by motion artifacts.

Thus far, our results relied on a measure of dynamic FC computed within a 45s sliding window. To examine if and how prediction accuracy varied with the size of the sliding window, we assessed the across-dataset model performance with dynamic FC computed with window sizes of 15s, 30s, 60s, and 75s. Across all window sizes, across-dataset prediction accuracy of the model was significantly above chance (**Fig. S2**, all *p* < .05), indicating that our results were robust to different window sizes. Altogether, these analyses suggest that dynamic FC encode moment-to-moment fluctuations in arousal and demonstrate that our results are robust to analytical decisions and low-level confounds.

### Arousal is encoded in FC patterns within and between large-scale functional networks

Is arousal encoded in the FCs within specific functional networks (e.g., dorsal attention network, default mode network), or is it encoded in the interactions between multiple networks? To answer this question, we examined the arousal network in each dataset. We term the FCs that were positively or negatively correlated with the arousal time course as positive or negative features respectively. The *Sherlock* arousal network includes 593 FC features (439 positive and 154 negative), and the *Friday Night Lights* arousal network includes 1578 FC features (848 positive and 730 negative).

We defined the set of overlap FC features between the *Sherlock* positive arousal network and the *Friday Night Lights* positive arousal network as the high-arousal network, and the set of overlap FC features between the *Sherlock* negative arousal network and the *Friday Night Lights* negative arousal network as the low-arousal network. We then assessed the degree of overlap in the high- or low-arousal networks across datasets. Both the high-arousal network (169 overlapping FC features, *p* < .001) and low-arousal network (68 overlapping FC features, *p* < .001) have an above-chance number of overlapping FC features across the two datasets. In contrast, the *Sherlock* high-arousal network does not significantly overlap with the *Friday Night Lights* low-arousal network (1 overlapping FC feature, *p* > .99), and the *Friday Night Lights* high-arousal network does not significantly overlap with the *Sherlock* low-arousal network (4 overlapping FC features, *p* > .99). Statistical significance of network overlap was assessed both using a hypergeometric cumulative density function (*53*) and using the Jaccard index as the measure of network similarity (*54*) (see Methods). The two methods generated consistent results.

To further understand how connections within and between large-scale functional networks constitute the high- and low-arousal network, we followed Yeo et al. (*49*) to group the 122 ROIs into 8 canonical functional networks, namely, the visual (VIS), somatosensory-motor (SOM), dorsal attention (DAN), ventral attention (VAN), limbic (LIM), frontoparietal control (FPN), default mode (DMN), and subcortical (SUB) networks. We asked whether particular functional networks were represented in the arousal network more frequently than chance (**Fig. 4A**). We computed the proportion of selected FCs among all possible FCs between each pair of functional networks, and assessed the significance of the proportions by comparing them against a null distribution of 10000 permutations where functional connections were randomly distributed across networks (see Methods). Connections between the DMN and FPN, DAN and FPN, and DAN and VAN, as well as connections within the DMN were positively associated with arousal. In contrast, connections between the DAN and VIS network were negatively associated with arousal (FDR-corrected *p* < .05). These results suggest generalizable neural representations of arousal are encoded within the DMN as well as between pairs of distributed networks.

**Fig. 4.**
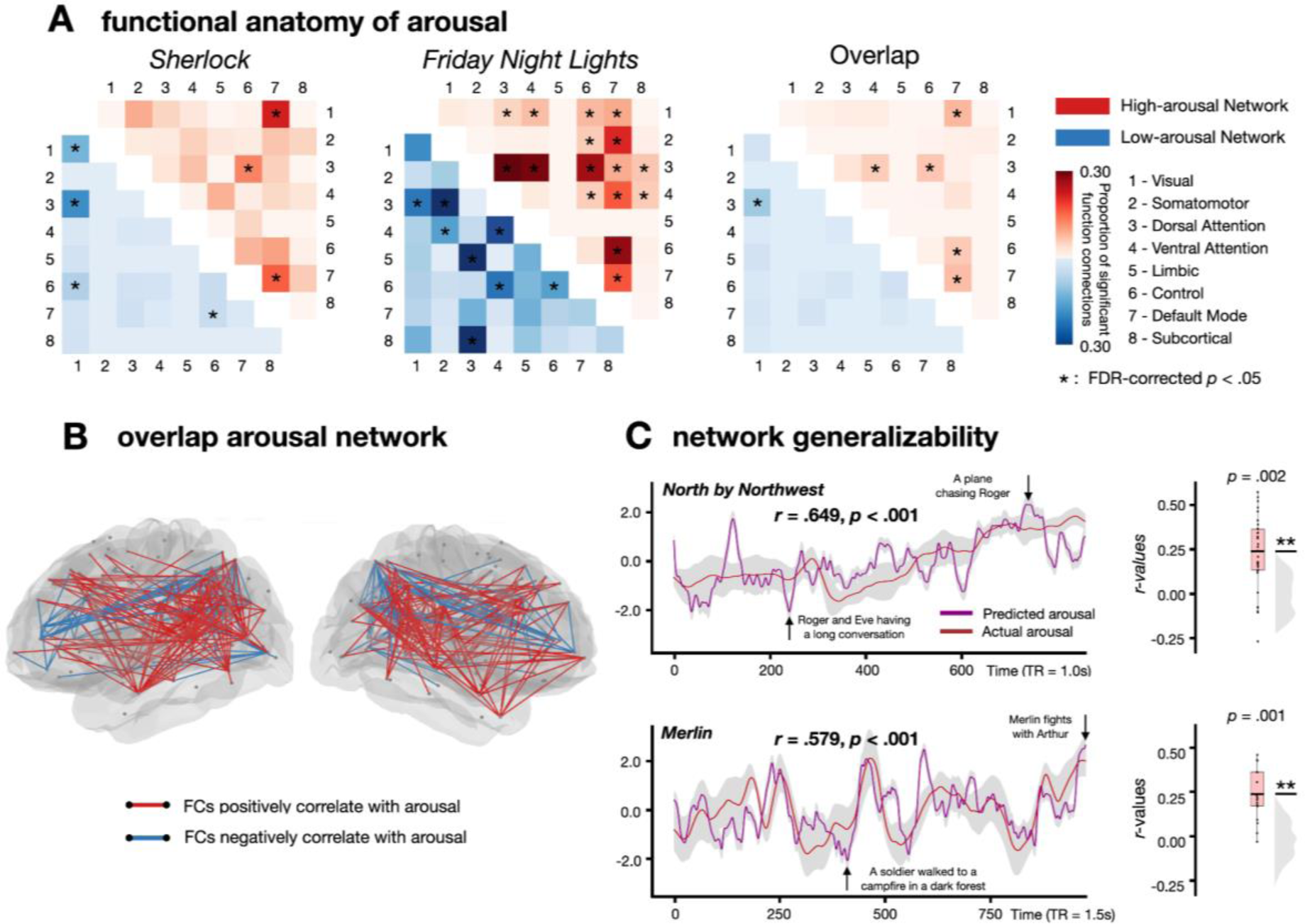
Functional anatomy of arousal. **A.** Arousal networks for *Sherlock*, *Friday Night Lights*, and their overlap. Each cell represents the proportion of selected FCs among all possible FCs between each pair of functional networks. The cells in the upper triangle represent the high-arousal network (red), and the cells in the lower triangle represent the low-arousal network (blue). Network pairs with above-chance selected FCs are indicated with an asterisk (one-tailed *t*-test, FDR-corrected *p* < .05). **B.** Visualization of the overlap arousal network. Each dot represents a node in the brain. The lines connecting two nodes show their functional connection. Red lines represent functional connections that positively correlate with arousal. Blue lines represent functional connections that negatively correlate with arousal. **C**. Connectome-based model trained on the overlap arousal network generalized to two more fMRI datasets, *Merlin* and *North by Northwest*. The left figure indicates how well the model could predict the group averaged experiences of arousal. We averaged the model predictions across participants watching the same movie and computed the correlation between the group-average model predictions and group-average arousal ratings. The right figure shows the predictive accuracy of the across-dataset prediction. The model was trained on a minimal set of FCs that predicted arousal in both *Sherlock* and *Friday Night Lights*, and tested separately on *Merlin* and *North by Northwest*. \*\**p* < .01, as assessed by comparing the empirical mean predictive accuracy against the null distribution.

A recent study analyzing the *Sherlock* dataset revealed that self-reported attentional engagement is correlated with scene-by-scene annotations of emotional arousal during narratives. This previous study characterized a set of functional networks that predicts engagement across datasets (*39*). The average engagement rating from that study was significantly correlated with the average arousal ratings that we collected (Pearson’s *r* = .778, *p* = .014) but not the average valence ratings (Pearson’s *r* = .085, *p* = .404). Relatedly, the arousal and engagement networks within the *Sherlock* dataset significantly overlap (361 overlapping FC features, *p* < .001; no overlap between the high arousal/low engagement and low arousal/high engagement networks). The shared FC features constitute 60.88% of FCs in the arousal network and 52.79% of FCs in the engagement network and were distributed across different functional networks (**Fig. S4**). However, predictions of a CPM trained on engagement ratings were not significantly correlated with behavioral arousal ratings (mean *r* = .498, *p* = .173; MSE = .754, *p* = .163; R^2^ = .245, *p* = .165). Moreover, the correlation was significantly lower than that of a model that was trained to predict behavioral arousal ratings (*r* = .498 vs. .558; paired-*t* test, *t*(15) = - 4.368, *p* < .001). In other words, even though engagement ratings were correlated with arousal ratings during the movie, a CPM trained to predict engagement ratings did not generalize to predict arousal. These results suggest that attentional engagement and arousal are related but non-synonymous constructs, with arousal possibly comprising an aspect of attentional engagement (*55*, *56*).

### Connectome-based models of arousal generalized to two more fMRI datasets

Our findings that CPMs trained on one movie predicted arousal in another largely distinct movie provide evidence that dynamic functional connectivity encodes generalizable representations of arousal. We then proceeded to test whether our identified arousal network would generalize to additional datasets. To that end, we analyzed two additional fMRI movie datasets with distinct genre, storyline, characters, and duration. The first was a 15-min clip from the movie *North by Northwest* (a spy thriller directed by Alfred Hitchcock; *N* = 32) collected by our group, and the second was a 25-min episode from BBC’s *Merlin* (a British fantasy adventure drama series; *N* = 18) from Zadbood and colleagues (*57*). To measure affective experience during the movie clips, we collected behavioral ratings of arousal and valence for both movies (*N* = 60, 15 in each condition). We tested the hypothesis that dynamic FC patterns encoded an arousal neural representation that generalizes across movie stimuli. To that end, we trained a CPM on both *Sherlock* and *Friday Night Lights* datasets with the overlap arousal network (169 positive FCs and 68 negative FCs; **Fig. 4B**) as input features. We then tested the model separately on the *North by Northwest* and *Merlin* datasets. Prediction accuracy was above-chance in both *North by Northwest* (mean *r* = .227, *p* = .002; MSE = .971, *p* < .001; R^2^ = .028, *p* = .003) and *Merlin* (mean *r* = .230, *p* = .004; MSE = .981, *p* < .001; R^2^ = .018, *p* = .010; **Fig. 4C**, right). Model-predicted arousal time courses also corresponded with the plot of each movie. For example, in *North by Northwest*, the most arousing moment predicted by the model occurred when the protagonist was being chased by a plane, while the least arousing moment occurred during a long conversation between characters (**Fig. S5**, top). Similarly, in *Merlin*, the most arousing moment predicted by the model occurred when the two main protagonists had a brawl in a tavern, while the least arousing moment occurred when a nameless soldier walked towards a campfire (**Fig. S5**, bottom). These results indicate robust out-of-sample generalizability, suggesting that the models were not merely capturing the idiosyncrasies of specific datasets. Instead, the arousal networks that we identified encode neural representations of arousal that generalized across diverse situations.

Given a movie-watching fMRI dataset, how well can the model predict the moment-to-moment arousal fluctuations of the average participant? In the prior analyses, we correlated the model predictions from each participant’s data with the group-average arousal ratings, and then averaged the correlations across the sample. This allowed us to assess the extent to which predictions from each individual matched the group average. If our goal was instead to maximize our ability to predict the group-average experience of arousal, we can first aggregate the model predictions across participants and predict average arousal ratings from the aggregated model predictions. To that end, we averaged the model predictions across participants watching the same movie and computed the correlation between the group-average model predictions and group-average arousal ratings. The group-average predicted arousal time course was significantly correlated with the group-average human-rated arousal time course in both *North by Northwest* (*r* = .649, *p* < .001) and *Merlin* (*r* = .579, *p* < .001; Fig. 4C, left). These results provide a measure of the model’s ability to predict arousal fluctuations of an average individual watching a movie that the model was not trained on, and suggest that our model could be used to generate moment-to-moment predictions of arousal during movie watching in new, independent datasets.

### Dynamic functional connectivity does not predict moment-to-moment valence

Having established that dynamic functional connectivity encodes emotional arousal, we examined whether the same approach can predict valence. Within-dataset CPMs did not show above-chance predictive accuracy in either *Sherlock* (mean *r* = .518, *p* = .193; MSE = .740, *p* = .195; R^2^ = .260, *p* = .194) or *Friday Night Lights* (mean *r* = .514, *p* = .094; MSE = .742, *p* = .090; R^2^ = .257, *p* = .091). Across-dataset predictive accuracy was also not significantly better than chance (train on *Sherlock* test on *Friday Night Lights*: mean *r* = -.041, *p* = .748; MSE = 1.204, *p* = .789; R^2^ = -.205, *p* = .791; train on *Friday Night Lights* test on *Sherlock*: mean *r* = -.016, *p* = .615; MSE = 1.177, *p* = .812; R^2^ = -.178, *p* = .820; **Fig. 3B**). Equivalence tests (*58–60*) indicated that these accuracies were statistically indistinguishable from zero within bounds defined by a minimal effect size of interest [-.100, .100] (see **Fig. S6**). One possible explanation of these null results is that the group-average valence ratings might not be sufficiently reliable across participants, potentially due to idiosyncratic experience of subjective valence. However, the inter-subject agreement of the valence ratings was not systematically different than that of arousal in both datasets and showed comparable overall inter-subject agreement across-datasets (valence: mean *r* = .677; arousal: mean *r* = .645; *t*-test: *t*(118) = .756, *p* = .451). These results suggest that the model’s poor performance at predicting valence fluctuations were unlikely to be due to lower reliability of the behavioral valence ratings. Another possible explanation is that the current specific window size we used in the predictive modeling did not capture the valence fluctuations on the right timescale. To rule out this possibility, we examined whether the valence model could generalize across datasets when the model was trained and tested on various different timescales. We used the same approaches as predicting arousal. None of the 25 combinations of window sizes showed above-chance accuracy in predicting valence across-dataset, suggesting that the null results were unlikely to be due to the choice of window size (**Fig. S3**).

The previous analyses assumed that valence is encoded as a single “bipolar” dimension, with positive and negative valence at opposing ends (*11*, *19*). An alternative possibility is that positive and negative valence are encoded separately in the brain (*61–64*), in which case, a CPM trained on data that spanned the entire valence spectrum would not perform well. To test this alternative possibility, we trained separate CPMs on positive and negative moments in each movie. For positive valence, we observed above-chance within-dataset predictive accuracy in *Friday Night Lights* (mean *r* = .762, *p* = .001; MSE = .459, *p* < .001; R^2^ = .541, *p* = .001) but not Sherlock (mean *r* = .431, *p* = .835; MSE = .823, *p* = .820; R^2^ = .176, *p* = .822). However, the model failed to generalize across-dataset (from *Sherlock* to *Friday Night Lights*: mean *r* = -.020, *p* = .528; MSE = 1.249, *p* = .565; R^2^ = -.251, *p* = .566; from *Friday Night Lights* to *Sherlock*: mean *r* = .088, *p* = .247; MSE = 1.059, *p* = .156; R^2^ = -.060, *p* = .157; Fig. S7B, left). For negative valence, neither within-dataset (*Sherlock*: mean *r* = .553, *p* = .400; MSE = .704, *p* =.386; R^2^ = .295, *p* = .383; *Friday Night Lights*: mean *r* = .647, *p* = .192; MSE = .607, *p* = .204 ; R^2^ = .391, *p* = .205) nor across-dataset (from *Sherlock* to *Friday Night Lights*: mean *r* = -.093, *p* = .875; MSE = 1.266, *p* = .958; R^2^ = -.268, *p* = .959; from *Friday Night Lights* to *Sherlock*: mean *r* = -.216, *p* = .898; MSE = 1.343, *p* = .885; R^2^ = -.345, *p* = .886; Fig. S7B, right) models showed significant predictive accuracy. As such, we were unable to identify generalizable neural representations of valence encoded in dynamic functional connectivity, even when considering positive valence and negative valence separately.

### Connectome-based models of valence failed to generalize even with more training data

One reason why the valence CPMs failed to generalize may be because they were trained on insufficiently diverse contexts. We thus sought to test if a valence CPM would generalize if trained on more data. To that end, we trained a CPM on the combined data from *Sherlock* and *Friday Night Lights*, with the 2067 FCs that significantly correlated with valence in the combined dataset as input features. This model failed to generalize to both *Merlin* (mean *r* = -.010, *p* = .611; MSE = 1.148, *p* = ,659; R^2^ = -.149, *p* = .660) and *North by Northwest* (mean *r* = -.140, *p* = .930; MSE = 1.200, *p* = .873; R^2^ = -.202, *p* = .874), indicating that despite training the model on multiple datasets, we were still unable to identify generalizable neural representations of valence encoded in dynamic functional connectivity.

### Univariate parametric mapping and multivariate pattern-based predictions of arousal and valence

Prior work has employed univariate parametric mapping and multivariate pattern analysis (MVPA) to examine the neural correlates of human affective experience (*26–30*). Here, we asked if we could identify generalizable neural representations of arousal and valence using these approaches.

We first examined whether the univariate parametric maps of arousal shared overlapping brain activations across movies. We implemented a generalized linear model (GLM) to examine the association between BOLD activity and arousal ratings. In *Friday Night Lights*, we found that the activation of voxels in regions across the whole brain, including the amygdala, insula, vmPFC, ventral striatum, anterior cingulate cortex (ACC) and orbitofrontal cortex (OFC), showed significantly positive correlations with arousal (TFCE-corrected *p* < .01). These results are consistent with the literature (*65–69*), suggesting the role of these regions in processing emotional salient stimuli. Similarly, in *Sherlock*, the activation in most of these regions showed positive correlations with arousal (**Fig. S8**) but did not survive controlling for multiple comparisons. We then performed a conjunction analysis (*70*) to identify the voxels consistently significantly related to arousal in both *Sherlock* and *Friday Night Lights* (TFCE-corrected *p* < .01). We observed clusters of voxels in regions including insula, ACC, and precuneus (**Fig. S9**).

Next, we ran the same GLM to test the association between BOLD activity and valence. We found that the activation in ACC and posterior cingulate cortex (PCC) was positively correlated with valence in both *Friday Night Lights* and *Sherlock* (**Fig. S8**). However, only the activation in *Friday Night Lights* survived controlling for multiple comparisons (p < .01). A conjunction analysis indicated that few voxels were significantly associated with valence in both movies (**Fig. S9**). These results suggest that, in our data, the same univariate activity encoded arousal, but not valence, across movies.

Having identified the voxels that were related to arousal and valence respectively, we proceeded to investigate whether the multivariate patterns of activation across voxels could predict the moment-to-moment fluctuations of arousal and valence. We used the same feature selection process and predictive modeling approach that we used for training the dynamic CPMs. Specifically, we applied a leave-one-subject-out cross-validation approach, where we examined all possible voxels across the whole brain and selected the voxels whose activity significantly correlated with the behavioral time course in the training set of *N* - 1 participants (one-sample Student’s *t* test, *p* < .01). We trained a non-linear support vector regression model in the set of *N* - 1 participants to predict the behavioral time course from these selected voxels, and tested on the held-out participants. Multivariate-pattern based models (MPMs) yielded significant within-dataset predictions of arousal in *Friday Night Lights* (mean *r* = .498, *p* = .024; MSE = .827, *p* = .073; R^2^ = .172, *p* = .085) but not Sherlock (mean *r* = .409, *p* = .160; MSE = .840, *p* = .099; R^2^ = .160, *p* = .109; **Fig. S10**). Importantly, the MPMs on arousal failed to generalize across datasets, both when training on *Sherlock* and testing on *Friday Night Lights* (mean *r* = .024, *p* = .327; MSE = 1.090, *p* = .466; R^2^ = -.091, *p* = .470), as well as when training on *Friday Night Lights* and testing on *Sherlock* (mean *r* = .054, *p* = .133; MSE = 1.020, *p* = .005; R^2^ = -.020, *p* = .006; **Fig. 5A**, left).

**Fig. 5.**
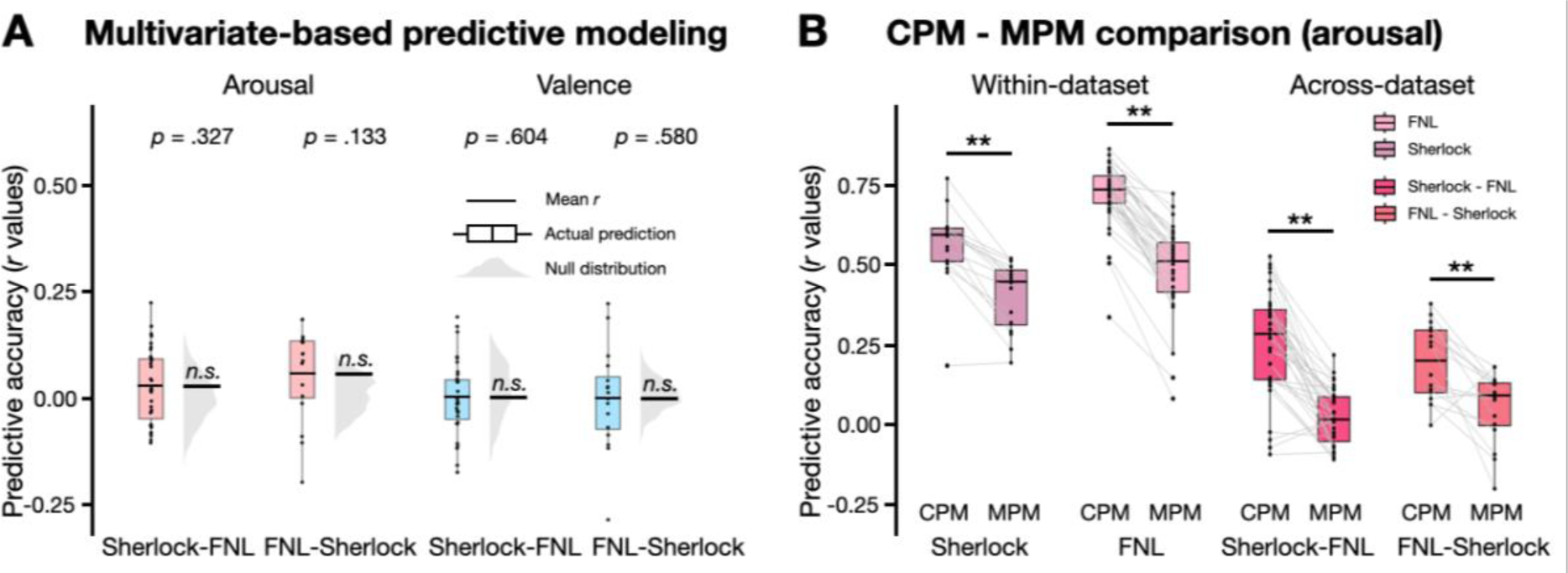
**A.** Multivariate-pattern based model performance in predicting arousal and valence across-datasets. n.s.: *p* > .05, as assessed by comparing the empirical mean predictive accuracy against the null distribution. **B.** Comparison between the predictive accuracy of connectome-based predictive modeling (CPM) and multivariate-based predictive modeling (MPM) for both within-dataset (left panel) and across-dataset predictions (right panel). Each datapoint in the box plot represents the predictive accuracy of each round of cross-validation. The gray lines connect the same held-out participant in CPM and MPM. \*\**p* < .01.

Model predictions of valence were not significant both within-dataset (*Sherlock*: mean *r* = .420, *p* = .120; MSE = .840, *p* = .099 ; R^2^ = .160, *p* = .099; *Friday Night lights*: mean *r* = .369, *p* = .344; MSE = .891, *p* = .385; R^2^ = .108, *p* = .406) and across datasets (from *Sherlock* to *Friday Night lights*: mean *r* = .001, *p* = .604; MSE = 1.108, *p* = .712; R^2^ = -.109, *p* = .718; from *Friday Night lights* to *Sherlock*: mean *r* = -.001, *p* = .580; MSE = 1.138, *p* = .913; R^2^ = -.139, *p* = .915 ; **Fig. 5A**, right). Together, these results indicate that we were not able to identify generalizable representations of either valence or arousal from multivariate activation patterns.

Finally, we investigated whether CPMs outperformed MPMs in predicting arousal by comparing their Fisher’s *z*-transformed predictive accuracies. CPM yielded significantly higher predictive accuracy than MPM both within-dataset (Paired *t*-test: *Sherlock*: *t*(15) = 6.963, *p* < .001; *Friday Night Lights*: *t*(34) = 18.056, *p* < .001) and across datasets (Paired *t*-test: from *Sherlock* to *Friday Night Lights*: *t*(34) = 8.512, *p* < .001; from *Friday Night Lights* to *Sherlock*: *t*(15) = 3.903, *p* = .001; **Fig. 5B**, right). These results suggest that dynamic functional connectivity patterns may encode a more reliable arousal representation than multivariate activation patterns.

## Discussion

This study utilized predictive modeling of affective experience to examine how valence and arousal states are represented in the human brain. We first showed that subjective behavioral ratings of valence and arousal were synchronized between individuals watching the same movie. We then identified a generalizable neural representation of emotional arousal across individuals and situational contexts encoded in the patterns of dynamic functional connectivity. Specifically, a model trained to predict arousal from dynamic functional connectivity during one movie generalized to a second movie, and the arousal representation further generalized to two additional datasets with different groups of participants. Moreover, predictive models trained on dynamic functional connectivity predicted arousal more accurately than predictive models trained on multivariate activation patterns, where the latter also did not generalize. In contrast to arousal, we found no evidence of a generalizable neural representation of valence. These results suggest that the neural encoding of arousal is intrinsically similar across different situations, while the neural encoding of valence might be specific to situations and individuals.

A strength of the current work is that, in addition to identifying the neural correlates of affective experience in a particular context, we explicitly tested the out-of-sample generalizability across multiple independent datasets with different stimuli. This is crucial because predictive models trained and tested within the same context may merely fit the idiosyncrasies of a specific stimulus. For instance, a model might learn the association between high arousal and the presence of a particular character in a narrative, but fail to predict arousal in a different narrative with different characters. A generalizable neural representation of a latent psychological construct should be consistent across various contexts and situations (*71–74*). Our results indicate that the neural representation of arousal encoded in dynamic functional connectivity meets this criterion of generalizability—the CPMs successfully predicted fluctuations in arousal in movies that the models had not previously encountered, which also differed in low-level audio-visual features, characters, plot, and genre. The robustness of the model across these stimuli suggests a shared neural representation of the subjective experience of arousal across different situations, highlighting an intrinsic similarity in how the brain encodes arousal from suspenseful moments in *Sherlock* to exciting moments in *Friday Night Lights*.

These findings align with and extend classic theories of emotion. In particular, the two-factor theory of emotion proposes that emotional experiences arise from the combination of physiological arousal and the cognitive interpretation of that arousal (*23*). To interpret arousal, individuals typically rely on contextual cues in the immediate environment. In cases where the source of arousal is ambiguous, people may mistakenly attribute their arousal to an incorrect source, a phenomenon known as the misattribution of arousal (*75–78*). For example, a person who arrives at a job interview after climbing up a flight of stairs might mistakenly attribute the arousal from the physical activity to nervousness about the interview, which can in turn affect their performance on the interview. In this view, arousal states are inherently confusable and susceptible to contextual misattribution. Our results suggest that such confusions may occur because the brain encodes arousal states similarly across different contexts. This shared representation of arousal could serve as a fundamental substrate upon which emotional experience is constructed and interpreted (*79–81*).

Our work builds upon earlier studies focusing on the neural correlates of arousal, which primarily explored the role of univariate activity and multivariate patterns (*26–30*, *82–84*). In our study, we found a multivariate activity pattern that encoded arousal within one dataset, but it failed to generalize when applied to a second dataset. Furthermore, a direct comparison revealed that connectivity-based models predicted arousal more accurately and more robustly across datasets than models based on multivariate activity patterns. These results suggest that while multivariate patterns capture arousal in specific contexts, connectivity patterns may tap into broader, stimulus-general neural mechanisms. In particular, the connectivity patterns may reflect the influence of global neuromodulatory systems, such as the norepinephrine system, which is known to regulate state-level changes across the brain and is crucial for modulating arousal in response to environmental demands (*85*).

A recent study by Young and colleagues (*86*) had found that connectivity within the salience network (corresponding closely to the ventral attention network), and between the salience network and the executive control network (corresponding closely to the frontoparietal control network) was associated with fluctuations in physiological arousal, as measured by heart rate. Our study complements and advances this prior work in several ways. In particular, we examined the neural representation of arousal in multiple movies with diverse affective contexts (e.g., excitement, suspenseful, comedic) and demonstrated that our model generalized across movies that the model was not trained on. In contrast to the Young et al. study, we identified connections within and between multiple networks, including the ventral attention network, the frontoparietal attention network, the dorsal attention network and the default mode network. Future work will be needed to disentangle whether these discrepancies are due to differences in the affective contexts tested (stress vs. more generalized arousal) or differences in how arousal was measured (physiological signal vs. subjective ratings).

The involvement of multiple large-scale functional networks in encoding arousal identified in our study aligns well with theories proposing that norepinephrine release during heightened arousal is associated with enhanced integration across functional networks in the brain (*87*, *88*). Network integration refers to a state of correlated activity across distinct functional networks and is thought to facilitate rapid and coordinated behavioral responses to environmental stimuli (*89*, *90*). For example, during heightened arousal, there may be increased coordination between regions involved in sensory processing, attention, and decision-making, reflecting a brain state optimized for processing salient information and preparing the body for action. As such, increased network integration induced by arousal could underlie the brain’s capacity to initiate a fight-or-flight response. The generalizability of the arousal representation identified in our study suggests that this arousal response may be context-independent rather than stimulus-specific.

Additionally, we discovered that connectivity-based arousal models trained on one group of participants generalized effectively to another group, suggesting that the neural representations of arousal are consistent not only across different stimuli but also among individuals. This indicates that there may be shared neural mechanisms underpinning the physiological and psychological aspects of arousal. Nonetheless, the presence of generalized representations does not rule out the coexistence of individual-specific patterns (*91*). Although our results underscore a common brain state linked with fluctuations in arousal, individual variations in neural connectivity may play an important role in how arousal is subjectively experienced and interpreted. Future studies could explore how unique neural patterns interact with the shared mechanisms identified here to contribute to the diversity of affective experiences.

Our research introduces a novel methodological approach for assessing moment-to-moment arousal ratings during movie watching. Collecting arousal ratings from human participants is labor-intensive and time-consuming. Given the increasing availability of open fMRI datasets that include movie-watching data (*92–94*), our model offers a valuable tool for researchers who wish to obtain continuous measures of arousal without the need for additional human ratings. A promising and important extension of the current work is to assess if the arousal representation would generalize to participants engaging in other tasks, such as viewing images (*28*) or making decisions (*95*). In contrast to arousal, we did not find a generalizable neural representation of valence. Notably, the one-to-average correlation in behavioral valence ratings comparable to that of the behavioral arousal ratings, suggesting that the subjective experience of overall valence was similarly synchronized across participants. Thus, the poor model performance was unlikely to be due to the idiosyncratic experience of subjective valence. An alternative explanation for the lack of generalizability of valence representations is that our models were trained across participants, while neural representations of valence may be individual specific. As we do not have data of the same participant watching multiple movies, this is not an explanation we were able to rule out. Isolating individual-specific neural representations of valence would likely require a “precision functional mapping” approach (*96*), where extensive functional data is collected from the same individual experiencing diverse affective contexts.

Our findings contrast with those of Chikazoe and colleagues (*32*), who identified a shared neural code for valence across different stimuli, modalities, and individuals. Our study is distinct from this earlier study in two key aspects. First, the earlier study utilized brief, context-free visual and gustatory stimuli, which may have constrained the subjective experience of valence. For instance, in the absence of context, viewing an image of a smiling baby may elicit a basic experience of pleasantness that would be similar to that when viewing an image of adorable puppies. In contrast, our study employed extended, narrative-driven stimuli, which likely evoke richer and more nuanced emotional experiences (*97*). Here, the positive feelings elicited by Sherlock’s witty banter with Watson in one scene may carry a different qualitative and neural signature compared to positive feelings felt during a victory by the football team in *Friday Night Lights*.

Second, participants in the earlier study provided ratings of subjective valence after each stimulus, which involved summarizing their affective experience into a numerical response on a common scale. This transformation could require a common neural representation, such that the valence of one stimulus could be compared to that of another (*98*). In contrast, participants in our study passively viewed the movies, which more closely resembles naturalistic affective experiences, where individuals do not typically have to quantify their emotions in real-time. If a common representation of valence is not computed spontaneously in the brain, it could explain why we were unable to detect a generalizable valence representation in our task.

Our results align with broader findings suggesting that valence may be encoded in a situation-specific rather than a generalizable manner (17, 96–98). For instance, a meta-analysis by Lindquist et al. (*99*) found no consistent brain regions associated with monotonically increasing or decreasing valence, suggesting that valence is encoded flexibly, contingent on the situational context. Altogether, our findings are consistent with theories of emotion positing that emotional experiences are not uniform across contexts but are heterogeneous and constructed from a mix of situational cues (*80*), cognitive appraisals (*21*), and social-environmental constraints (*100*). Two experiences, even if similarly positively valenced, engage different neural computations and circuits in different contexts. Thus, the neural representation of valence would not be similar across situations. As our study did not exhaustively rule out other possibilities of how a generalizable representation of valence might be encoded in the brain, this interpretation remains tentative. Nevertheless, we believe that explorations into the neural mechanisms that act on generalizable representations of arousal to construct situation-specific representations of valence would be a fruitful direction for future research.

In summary, our study identified a generalizable neural representation of arousal encoded in dynamic functional connectivity, but did not find a parallel generalizable representation of valence. The findings highlight the relationship between arousal states and the dynamics of large-scale functional networks. This work extends our understanding of how affective experience is encoded in the brain, and provides a methodological approach of probing the neural basis of affective experience in naturalistic contexts using functional neuroimaging and machine learning techniques. Our study paves the way for future studies investigating the relationship between everyday affective states from neural dynamics. By integrating our approach with deep learning methods that can extract stimulus features from multimodal sensory stimuli (*101*), future studies can explore the mechanisms by which specific aspects of a stimulus drive arousal responses and influence connectivity patterns.

## Materials and Methods

### Subjects

One hundred and twenty individuals participated in the behavioral experiment (83 female, mean age 20.45 ± 2.15 years). Subjects were randomly assigned to four experimental conditions, namely, *Sherlock* valence, *Friday Night Lights* valence, *Sherlock* arousal, and *Friday Night Lights* arousal. Each condition includes data from 30 subjects. The rating time courses were significantly correlated across individuals in each condition (**Fig. S1**). Participants provided informed written consent before the start of the study in accordance with the experimental procedures approved by the Institutional Review Board of the University of Chicago and were compensated for participation with cash or credits for class.

### Stimuli and experiment design

A 48min 6s segment of the BBC television series *Sherlock* (*44*), and a 45min 18s segment of the NBC serial drama *Friday Night Lights* (FNL) (*45*) were used as the audio-visual movie stimuli in the study. The 30s audiovisual cartoon (*Let’s All Go to the Lobby*) was removed from the original *Sherlock* movie stimuli. Both stimuli were divided into two segments (23min and 25min 6s for *Sherlock*, 23min 45s and 21min 33s for *FNL*). We prepended a 5s countdown video to the beginning of each segment to prepare participants to the beginning of the movie. Before rating the movie, participants performed a practice task where they rated valence or arousal of a short video from the movie *Merlin* (*57*) (2:03-3:10 for valence, and 5:13-6:19 for arousal) to get familiar with the task.

Participants watched either *Sherlock* or *FNL*, and continuously rated either how positive or negative (valence conditions), or how aroused or not aroused (arousal conditions) the videos made them feel at each moment while watching the video. Participants who provided valence ratings were told that positive valence refers to when they were feeling pleasant, happy, excited, and negative valence refers to when they were feeling unpleasant, sad, angry. Participants who provided arousal ratings were told that high arousal refers to when they were feeling mentally or physically alert, activated, and/or energized, and low arousal refers to when they were feeling very mentally or physically slow, still, and/or de-energized. Participants pressed “left” or “right” keys on a keyboard to adjust a slider bar on a 1-25 scale. The two ends of the scale were labeled as “Very Negative” and “Very Positive” for participants in the valence conditions, and “Completely Not Aroused” to “Completely Aroused” for participants in the arousal conditions. Participants were encouraged to move the slider bar for even slight changes in how they felt. The button presses were recorded by a program coded in jsPsych (www.jspsych.org). Participants had the option to take a break after they watched the first video segment and continued to watch the second segment whenever they felt ready to do so. Participants completed a post-experiment survey after they finished the main rating experiment, where they reported the overall valence or arousal rating of the two segments, demographics and whether they had watched the video before. 15, 17, 3, and 5 participants had viewed the movie episode before the experiment in the *Sherlock*-valence, *Sherlock*-arousal, *FNL*-valence, *FNL*-arousal condition respectively. As approximately half of the participants had previously watched *Sherlock*, we checked if the reliability of the ratings differed between participants who had or had not previously watched the episode. There was no significant difference in the pairwise rating similarity of the two groups both for the arousal (*t*(29) = 1.157, *p* = .257) and valence ratings (*t*(29) = 1.543, *p* = .134). In addition, the arousal models trained on the behavioral ratings from participants who had (mean *r* = .234, p = .019; MSE = .990, *p* = .024 ; R^2^ = .009, *p* = .025) and had not (mean *r* = .297, p = .008; MSE = .929, *p* = .004 ; R^2^ = .070, *p* = .005) seen the movie generalized to *Friday Night Lights*.

### Movie analysis and annotations

We extracted 7 low-level visual features (hue, saturation, pixel intensity, motion energy, presence of a face, whether the scene was indoor or outdoor, presence of written text) and 3 low-level auditory features (amplitude, pitch, presence of music) from both *Sherlock* and *FNL*. We computed hue, saturation, pixel intensity (‘rgb2hsv’), audio amplitude (‘audioread’) and pitch (‘pitch’) in MATLAB (*51*) (R2022b, The Mathworks, Natick, MA), and motion energy (pliers.extractors. FarnebackOpticalFlowExtractor) in Python (*102*) (version 3.9). Additionally, author J.K. coded whether each video frame contained a face (presence of face = 1, absence of face = 0), written text (presence of written text = 1, absence of written text = 0), whether the scene was indoor or outdoor (indoor = 1, outdoor = 0), and whether music played in the background (presence of music = 1, absence of music = 0).

### Behavioral data analysis

Considering that rating changes can happen at any time during the movie and the time points of the changes differ across subjects, we resampled the ratings to one rating per TR (1.5s for *Sherlock* and 2s for *FNL*). Resampled time courses of the two movie segments from the same subject were concatenated, and z-scored across time. To compute the intersubject correlation of the affect ratings, we computed the Pearson correlation between each individual’s subjective rating timecourse to the group-averaged rating timecourse with this individual left out. We calculated the similarity within each condition (i.e., *Sherlock*-valence, *Sherlock*-arousal, *FNL*-valence, *FNL*-arousal). The pairwise *r*-values were then Fisher *z*-transformed and averaged across individuals. An inverse transform was then applied to the resulting average *z*-value to obtain the average mean intersubject correlation for a given condition. To test whether group-mean ISC significantly deviated above zero, we applied a nonparametric permutation approach (10000 permutations) where 50% chance of sign-flipping was applied to every subject’s rating similarity before averaging into group-mean. The normalized rating time courses of all subjects within each condition were averaged, and the group-average time courses were treated as the ground-truth affective experience time-locked to the stimuli.

Further fMRI analysis in the study involved the application of a tapered sliding window. In order to align the behavioral time courses to fMRI data time courses, we convolved the group-average time courses with the hemodynamic response function (HRF), and applied a tapered sliding window to the convolved behavioral time courses. We applied a sliding window size of 30 TR (45s) for *Sherlock* and 23 TR (46s) for *FNL*, and a step size of 1TR and a Gaussian kernel σ = 3TR. We followed Song et al. (*39*) in determining the weights for cropping the tail of the Gaussian kernel at the beginning and end of the time courses.

### MRI acquisition and preprocessing

We acquired the raw structural and functional images of the *Sherlock* dataset from Chen et al. (*44*), and the *FNL* dataset from Chang et al. (*45*). The *Sherlock* data were collected on a 3T full-body scanner (Siemens Skyra) with a 20-channel head coil. Functional images were acquired using a T2*-weighted echo-planar imaging (EPI) pulse sequence (TR/TE = 1500/28 ms, flip angle 64°, whole-brain coverage 27 slices of 4 mm thickness, in-plane resolution 3 × 3 mm^2^, FOV 192 × 192 mm^2^), with ascending interleaved acquisition. Anatomical images were acquired using a T1-weighted MPRAGE pulse sequence (0.89 mm^3^ resolution). The *FNL* data were collected on a 3T Philips Achieva Intera scanner with a 32-channel head coil. Functional images were acquired in an interleaved fashion using gradient-echo echo-planar imaging with prescan normalization, fat suppression and an in-plane acceleration factor of two (i.e., GRAPPA 2), and no multiband (i.e., simultaneous multislice) acceleration (TR/TE = 2000/25 ms, flip angle 75°, resolution 3 mm^3^ isotropic voxels, matrix size 80 by 80, and FOV 240 × 240 mm^2^). Anatomical images were acquired using a T1-weighted MPRAGE sequence (0.90 mm^3^ resolution).

We applied the same preprocessing pipeline to the *Sherlock* and *FNL* dataset, which included MCFLIRT motion correction, high-pass filtering of the data with a 100-ms cut-off, and spatial smoothing using a Gaussian kernel with a full-width at half-maximum (FWHM) at 5 mm. The functional images were resampled to 3 mm^3^ isotropic space, and registered to participants’ anatomical images (6 d.f.), then to the Montreal Neurological Institute (MNI) space using affine transformation (12 d.f.). Preprocessing was performed using FSL/FEAT v6.00 (FMRIB software library, FMRIB, Oxford, UK).

### Dynamic connectome-based predictive modeling

We applied dynamic connectome-based predictive modeling, an approach introduced in Song et al. (*39*) and available at https://github.com/hyssong/NarrativeEngagement, to predict arousal and valence. We first extracted BOLD signals across the whole brain using the 114-ROI cortical parcellation scheme of Yeo et al., along with the 8-ROI subcortical parcellation from the Brainnetome atlas (bilateral amygdala, hippocampus, basal ganglia, and thalamus), yielding a total of 122 ROIs. We averaged the blood-oxygen-level dependent (BOLD) time courses of all voxels in each ROI. For each dataset, the dynamic functional connectivity (FC) patterns were extracted from the ROI-based BOLD time courses using a tapered sliding window approach, where the Fisher’s z-transformed Pearson’s correlation between the BOLD time courses of every pair of ROIs were computed within each tapered sliding window (Window size - *Sherlock*: 30 TRs = 45s; *FNL*: 23 TRs = 46s). Hyperparameters of the sliding window for brain data were the same as those for behavioral data (Step size: 1TR, Gaussian kernel σ = 3TR). In the complementary analysis where we examined how the predictive accuracy of the model would change over different timescales of dynamic functional connectivity patterns, we also used window sizes of 15s, 30s, 60s and 75s in addition to the original 45s or 46s in the main analysis. The same window sizes were also applied to the behavioral ratings to match the timescale of function connectivity patterns.

#### Within-dataset FC prediction

Within each dataset, we built separate CPMs to predict valence or arousal time courses using leave-one-subject-out cross-validation. In each round of cross-validation, we conducted feature selection, where functional connections (FC) that significantly correlated with affect time course in the training set of *N* - 1 participants were selected (one-sample Student’s *t* test, *p* < .01). A non-linear support vector regression model (python package svm.SVR from sklearn, https://scikit-learn.org/, kernel = “rbf”) was trained in the set of *N* - 1 participants to predict the group-average behavioral time course from the selected FC features, and tested on the held-out participant. The predictive accuracy of each round of cross-validation was calculated as the Pearson’s correlation between the predicted and ground-truth arousal time course. Correlation coefficients of all rounds of cross-validation were Fisher’s z-transformed, averaged across all cross-validation folds, and transformed back to an average r-value as the final measure of model performance. To assess statistical significance, we compared model performance against null distributions generated by training and testing the model on phase-randomized behavioral ratings (1000 permutations). We assumed a one-tailed significance test, with *p* = (1 + number of null *r* values ≥ empirical *r*)/(1 + number of permutations).

#### Across-dataset FC prediction

The set of FCs selected in every round of cross-validation in within-dataset prediction was used as the features in across-dataset prediction. All subjects’ data in the training dataset (e.g., *Sherlock*) were used to train a non-linear SVM to learn the mapping between the patterns of selected FC and the group-average affect time course. The trained model was applied to the held-out dataset, to predict each subject’s valence or arousal time course from their FC patterns. The predictive accuracy and statistical significance were evaluated in the same way as within-dataset FC prediction.

### Arousal networks

For each dataset, we termed the set of functional connections that were selected in every round of cross-validation in predicting arousal as that dataset’s arousal network. We defined FC features that positively, or negatively predict arousal as the high- or low-arousal networks, respectively. To examine the relationship between the arousal network and canonical macroscale functional networks in the brain, we grouped the 122 ROIs into 8 functional networks: the visual (VIS), somatosensory-motor (SOM), dorsal attention (DAN), ventral attention (VAN), limbic (LIM), frontoparietal control (FPN), default mode (DMN), and subcortical (SUB) networks. These networks were previously identified based on resting-state functional connectivity MRI of 1000 subjects (*49*). The set of functional connections that occurred in both the *Sherlock* and *FNL* arousal networks were defined as the overlapping arousal network. We used two methods to calculate statistical significance of network overlap. One was using the hypergeometric cumulative density function, which returns the probability of drawing up to x of K possible items in n drawings without replacement from an M-item population (*53*). This was implemented in MATLAB as: p = 1 – hygecdf(x, M, K, n), where *x* refers to the number of overlapping edges, K and n refers to the number of edges in each of the two networks, and M refers to the total number of edges (7381). The other was the Jaccard index (*54*), which measures the similarity between two sets by computing the ratio between the size of their intersection and the size of their union. It was implemented using sklearn.metrix.jaccard_score (https://scikit-learn.org/stable/modules/generated/sklearn.metrics.jaccard_score.html). To test the significance of the jaccard score, we compared the actual Jaccard score with a null distribution, generated by shuffling the position of selected FCs in the brain network (keeping the same number of selected FCs) before calculating the Jaccard score. These two methods generated consistent results.

We calculated the proportion of selected FCs relative to the total number of possible connections between each pair of functional networks. To statistically assess whether particular functional networks are represented in the arousal network more frequently than chance, we computed the proportion of selected FCs among all possible FCs between each pair of functional networks. To assess the significance of these proportions, we compared the proportions with a null distribution of 10000 permutations. We generated the null distribution by the following steps. First, we shuffled the positions of selected FCs (e.g., 593 for *Sherlock* high-arousal network) in the whole FC networks (i.e., 7381 FCs). Second, we divided the FC networks into the 8 canonical macroscale functional networks. Third, we computed the proportions for every pair of functional networks following the same procedure. Finally, the significance of each proportion is computed by comparing the actual value with the null distribution of 10000 permutations using a one-tailed t-test (fdr-corrected).

### Engagement behavioral ratings and networks

We acquired the engagement behavioral ratings of the *Sherlock* movie from Song et al., (*39*) GitHub repository (https://github.com/hyssong/NarrativeEngagement). We averaged individual rating time courses across subjects, cropped the 52 TRs of the two cartoon segments (1-26 TR, 947-972 TR) from the group-average engagement rating time course, and z-normalized the rating time course. We computed the Pearson correlation between the average engagement and arousal rating time courses. The significance of the correlation was assessed by comparing the actual correlation with a null distribution, generated by circular shifting the average engagement rating time course 1000 times.

We trained a CPM on the engagement time course, and used the CPM to predict the arousal time course following a leave-one-subject-out cross validation procedure. The engagement time course was smoothed with the same tapered sliding window size as the arousal time course and functional connectivity patterns. The predictive accuracy in each run of cross-validation was calculated as the Pearson’s correlation between model predictions and the ground-truth arousal time course. We then performed a paired *t*-test to test if model accuracy in predicting arousal was different between the CPM trained on engagement and the CPM trained on arousal.

To examine whether the neural representations of engagement and arousal shared underlying functional networks, we compared our arousal network with the engagement network acquired from Song et al. (*39*). This previous work used leave-one-subject-out cross-validation to predict attentional engagement from dynamic functional connectivity in the *Sherlock* dataset. The *Sherlock* engagement FC network was the set of FCs selected in every round of cross-validation in predicting engagement. We computed the arousal-engagement overlap network by taking the overlapping FCs between the *Sherlock* arousal FC network (439 positive and 154 negative FC features) and the *Sherlock* engagement *FC* network (583 positive and 102 negative FC features), resulting in a network with 287 positive and 74 negative FC features.

### Overlap arousal network predictions on two more fMRI datasets

To examine whether the overlap network between the *Sherlock* and *Friday Night Lights* arousal networks encoded a generalizable arousal representation, we trained a model on the overlap arousal network and tested on two more fMRI datasets, *North by Northwest* and *Merlin*.

The movie stimuli of the *North by Northwest* dataset was a 14:49-min long segment of the movie *North by Northwest*, an American spy thriller directed by Alfred Hitchcock. The fMRI data were collected on a 3T Philips Ingenia scanner at the MRI Research Center at the University of Chicago as part of an ongoing two-session study collecting task, movie, and annotated rest data. Only the data where participants watched the *North by Northwest* clip were analyzed here. Structural images were acquired using a high-resolution T1-weighted MPRAGE sequence (1.000 mm^3^ resolution). Functional BOLD images were collected on a 3T Philips Ingenia scanner with a 32-channel head coil (TR/TE = 1000/28 ms, flip angle 62°, whole-brain coverage 27 slices of 3 mm thickness, in-plane resolution 2.5 × 2.5 mm^2^, matrix size 80 by 80, and FOV 202 × 202 mm^2^). Functional scans were acquired during the movie segment in a single continuous run (951 TRs). The functional run included an extra 57 TRs after the video ends where participants were instructed to stare at the cross over a blank screen. These TRs were cropped from the end of each functional image to match the length of the movie stimuli. Data from 32 participants were included here. The movie stimuli of the *Merlin* dataset was a 25-min long episode from BBC’s *Merlin* (a British fantasy adventure drama series). The fMRI data was acquired from openneuro. The fMRI preprocessing pipeline and dynamic functional connectivity analysis were the same as those of *Sherlock* and *Friday Night Lights*. Additional behavioral ratings on valence and arousal were collected on both *North by Northwest* and *Merlin* movie clips. The continuous rating paradigm was implemented in a single continuous run, separately for *North by Northwest* and *Merlin*. The same behavioral preprocessing and analysis pipeline as those of *Sherlock* and *Friday Night Lights* were performed for *North by Northwest* and *Merlin*.

The set of FCs in the overlap arousal network between the *Sherlock* arousal network and the *Friday Night Lights* arousal network was used as the input features in the current across-dataset prediction to test its generalizability. The data from both all subjects in *Sherlock* and all subjects in *Friday Night Lights* was used to train a non-linear SVM to learn the mapping between the patterns of selected FC and the group-average arousal time course. The trained model was applied separately to the *North by Northwest* and *Merlin* dataset, to predict each subject’s arousal time course from their FC patterns. The predictive accuracy and statistical significance were evaluated in the same way as within-dataset FC prediction.

### Examining out-of-sample generalizability of valence predictions with additional training data

To examine whether a CPM of valence could generalize with more training data, we trained a CPM on the combined data from *Sherlock* and *Friday Night Lights*. In the combined neural data, we selected the 2067 FCs that significantly correlated with valence as input features. We trained a non-linear support vector regression model to predict the valence time courses from the combined neural data of the selected FCs, and tested this trained model separately in the *North by Northwest* and *Merlin* datasets.

### Defining positivity and negativity

To identify moments with positive valence (positivity), we took the raw behavioral valence ratings (range 1 - 25, 13 = neutral), averaged across participants rating the same movie, extracted the time segments where the group-averaged raw behavioral rating was above 13, and concatenated them into one single time course. We *z*-normalized and convolved the new rating time course with a HRF, and smoothed the new rating time course using a tapered sliding window (window size *Sherlock*: 30 TRs; *FNL*: 23 TRs). Moments with negative valence (negativity) were identified following the same procedure except that ratings that were below 13 were extracted from the group-average valence rating time course. The *Sherlock* movie had 1114 and 800 TRs of positivity and negativity respectively, and the *FNL* movie has 758 and 594 TRs of positivity and negativity respectively (**Fig. S5**).

To build predictive models to predict positivity and negativity from brain fMRI activity, we preprocessed the brain data to assess how they would be associated with the positivity and negativity rating time courses. We separately extracted the TRs corresponding to the behavioral positivity and negativity moments from the 122 parcels’ activation magnitude time courses, and applied the same tapered sliding window approach as described above to extract the dynamic functional connectivity patterns.

### Univariate generalized linear model

We ran generalized linear models (GLM) to identify the voxels whose BOLD activity were associated with valence or arousal in each movie. A Separate model was built for each of the four experimental conditions: *Sherlock*-valence, *FNL*-valence, *Sherlock*-arousal, and *FNL*-arousal. The GLMs were estimated throughout the whole brain using FSL/FEAT v.6.00.. Correction for multiple comparisons was performed using threshold-free cluster enhancement (TFCE correction) with an alpha level of 0.05, as implemented by the randomise tool in FSL (*103*). To assess the overlap in statistical maps across movies, we performed a minimum statistic test compared to the global null (MS/GN) conjunction analysis (*70*) to identify the voxels in the brain whose activations were consistent across Sherlock and FSL (*p* < .01). The test was run separately for arousal and valence.

### Multivariate-pattern based predictive modeling

To examine whether the multivariate patterns of activation encode valence and arousal, we employed the same analysis pipeline as that of dynamic CPM, except that the input features of the models were now the time courses of each voxel (Tapered sliding window size - *Sherlock*: 30 TRs = 45s; *FNL*: 23 TRs = 46s) following the approach described in Song et al.(*39*). Hyperparameters of the sliding window for brain data were the same as those for behavioral data (Step size: 1TR, Gaussian kernel σ = 3TR).

#### Within-dataset prediction

Within each dataset, we built separate multivariate pattern-based predictive models (MPM) to separately predict valence or arousal time courses using a leave-one-subject-out cross-validation approach. In each round of cross-validation, we conducted feature selection, where voxels whose activity significantly correlated with the behavioral time course in the training set of *N* - 1 participants were selected (one-sample Student’s *t* test, *p* < .01). The remaining procedures were identical to how we trained and tested the CPMs. The large number of voxels selected (24333 features) for each iteration posed a computational challenge for the permutation tests. To reduce computational cost, we ran 100 instead of 1000 permutations. We assumed a one-tailed significance test, with *p* = (1 + number of null *r* values ≥ empirical *r*)/(1 + number of permutations).

#### Across-dataset prediction

The set of voxels selected in every round of cross-validation in within-dataset prediction was used as the features in across-dataset prediction. All subjects’ data in the training dataset (e.g., *Sherlock*) were used to train a non-linear SVM to learn the mapping between the patterns of selected voxels and the group-average affect time course. The trained model was applied to the held-out dataset, to predict each subject’s affect time course from their multivariate patterns. The predictive accuracy and statistical significance were evaluated in the same way as within-dataset predictions, except that the number of permutations to generate the null distribution was 1000.

## Acknowledgments

We thank Janice Chen and colleagues for open sourcing the *Sherlock* dataset, Luke Chang and colleagues for open sourcing the *Friday Night Lights* dataset, and Asieh Zadbood and colleagues for open sourcing the *Merlin* dataset. We thank the MRI Research Center at the University of Chicago (RRID: SCR_024723) for assisting with the collection of the *North by Northwest* dataset. We thank Emily Finn for sharing code on calculating optical flow. We thank Yizhou (Louisa) Lyu for collecting part of the behavioral rating data. We thank members of the Bainbridge, Leong, Rosenberg, Bakkour (BLRB) community, especially members of the Motivation and Cognition Neuroscience (MCN) Lab and Cognition, Attention, and Brain (CAB) Lab at the University of Chicago for their helpful feedback. Our work was supported by the University of Chicago Social Sciences Division, the University of Chicago Neuroscience Institute, and resources provided by the University of Chicago Research Computing Center.

## Funding

National Science Foundation BCS-2043740 (M.D.R.)

## Author contributions

Conceptualization: J.K., H.S., Z.B., M.D.R., Y.C.L

Methodology: J.K., H.S., M.D.R., Y.C.L.

Data collection: J.K., Z.B.

Data curation, Formal analysis, & Visualization: J.K.

Supervision: M.D.R., Y.C.L.

Funding acquisition: M.D.R.

Writing—original draft: J.K., Y.C.L.

Writing—review & editing: J.K., H.S., M.D.R., Y.C.L.

All the authors approved the final manuscript for submission.

## Competing interests

Authors declare that they have no competing interests.

## Data and materials availability

All data are available in the main text or the supplementary materials.

The behavioral and brain data, the analysis code, as well as a step-by-step instruction to run the analysis code, are openly available on GitHub: https://github.com/jinke828/AffectPrediction/

## Notes

### Competing Interest Statement

The authors have declared no competing interest.

### Summary of Updates

Added complementary analysis; Supplemental files updated

