## Supplement for "Dynamic functional connectivity encodes generalizable representations of emotional arousal across individuals and situational contexts"

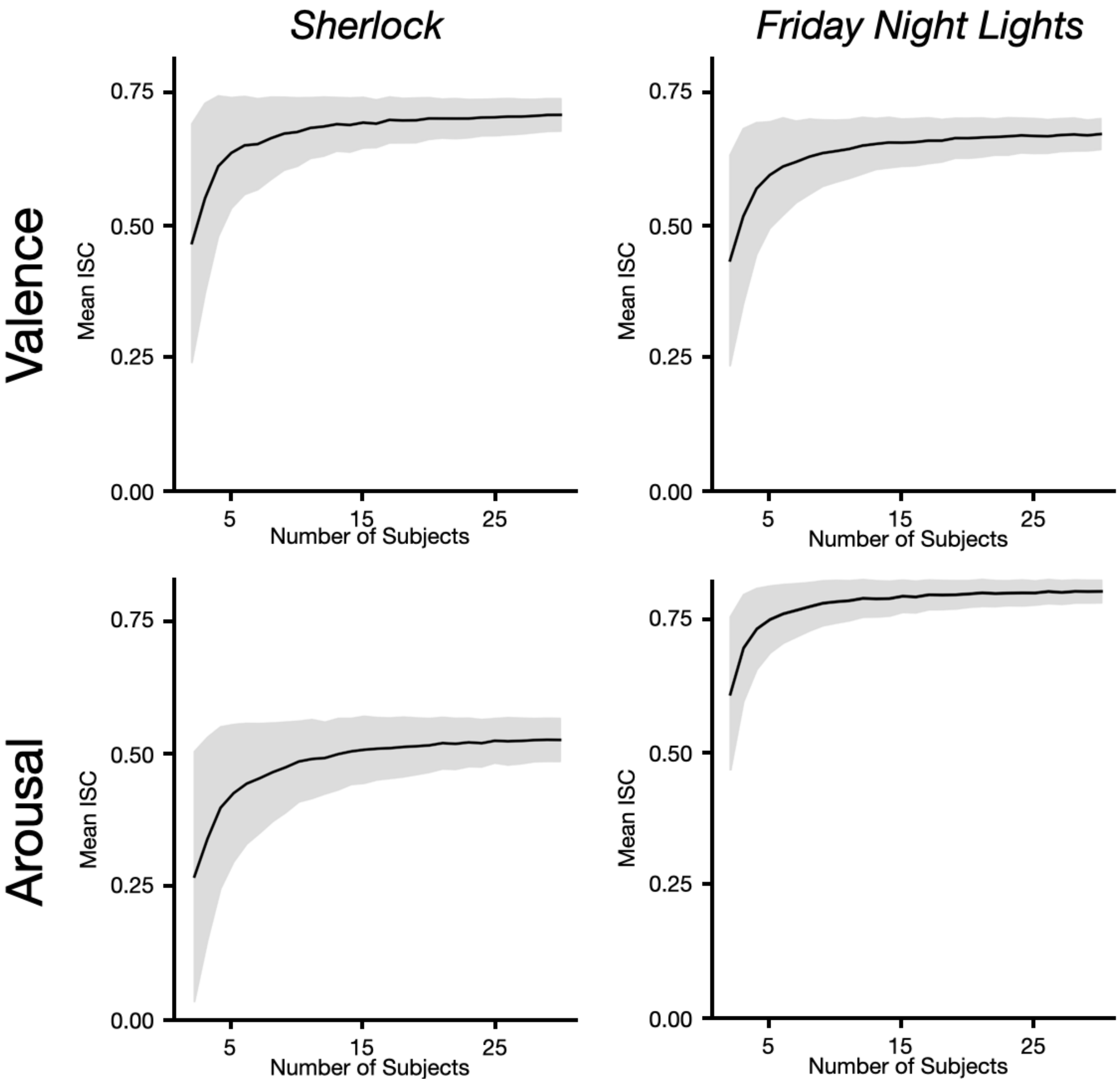


**Supplementary Figure 1. Leave-one-participant-out rating similarity as a function of the number of subjects**. The x-axis represents the number of subjects. In each condition, the number of subjects, *k*, increased from 2 to 30. Corresponding to the point *k* on the x-axis, *k* subjects were randomly selected from all subjects for 1000 times with replacement, where each time the Fisher's z-transformed group-average ISC was computed. The gray area represents the standard deviation of the distribution of permutations. The group-average rating similarity stabilizes as the number of subjects increases, suggesting it is unlikely for the group-average behavioral rating to increase in precision with more subjects.

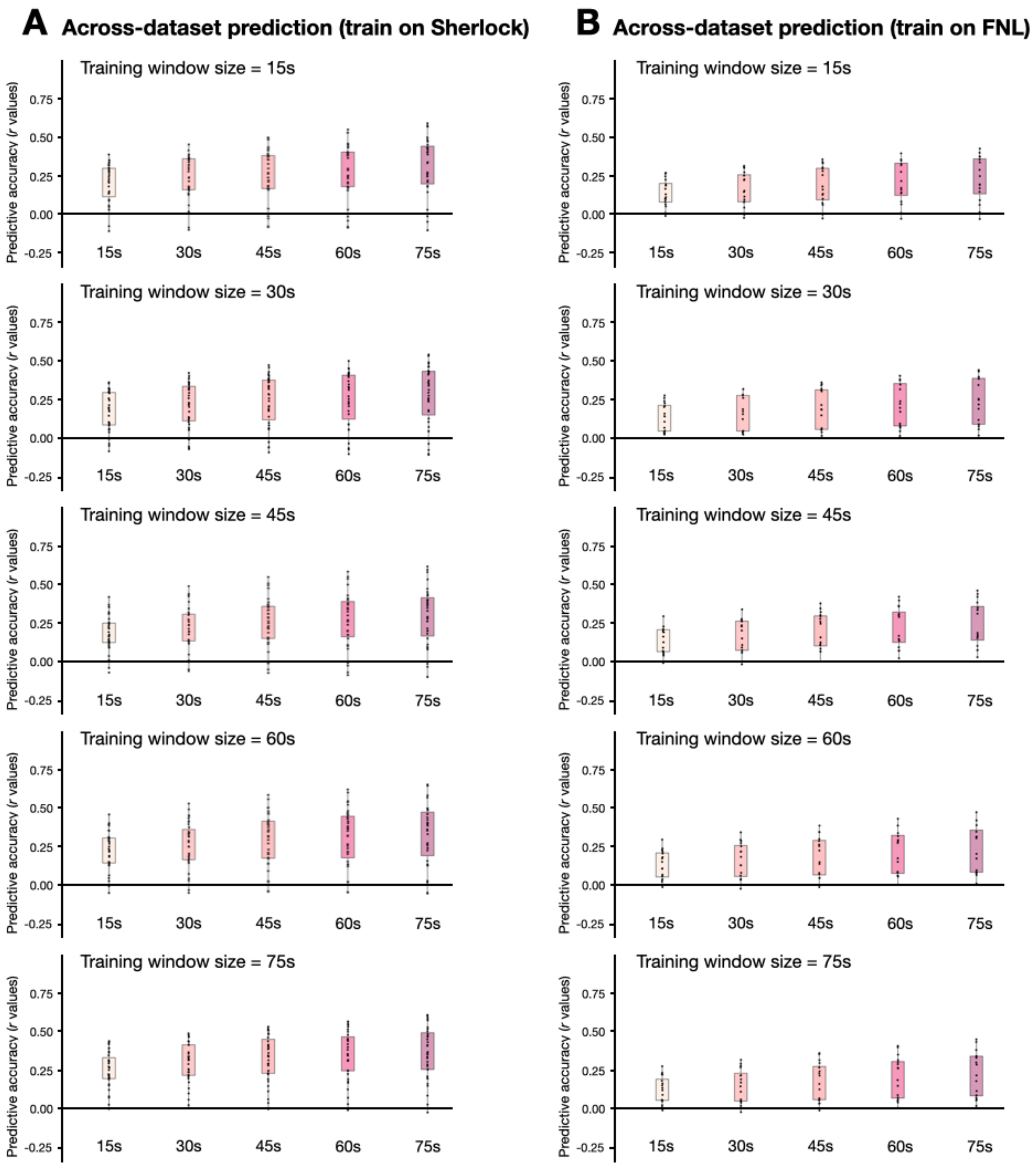


**Supplementary Figure 2. Across-dataset model predictive accuracy on arousal as a function of sliding window size**, when training on *Sherlock* and testing on *Friday Night Lights* **(A)** and when training on *Friday Night Lights* and testing on *Sherlock* **(B)**. We extracted dynamic functional connectivity patterns from both *Sherlock* and *Friday Night Lights* using 5 different sizes of tapered sliding window: 15s, 30s, 45s (the original size in the main analysis), 60s and 75s, and tested each model on test data at each window size, resulting in 50 conditions (2 datasets x 5 window sizes at training x 5 window sizes at testing). For example, in the top figure on the left, we trained the model in *Sherlock* with window size of 15s, and tested on *Friday Night Lights*, with window sizes of 15s, 30s, 45s, 60s and 75s. All 50 conditions of the across-dataset predictions on arousal showed significantly higher accuracy than chance.

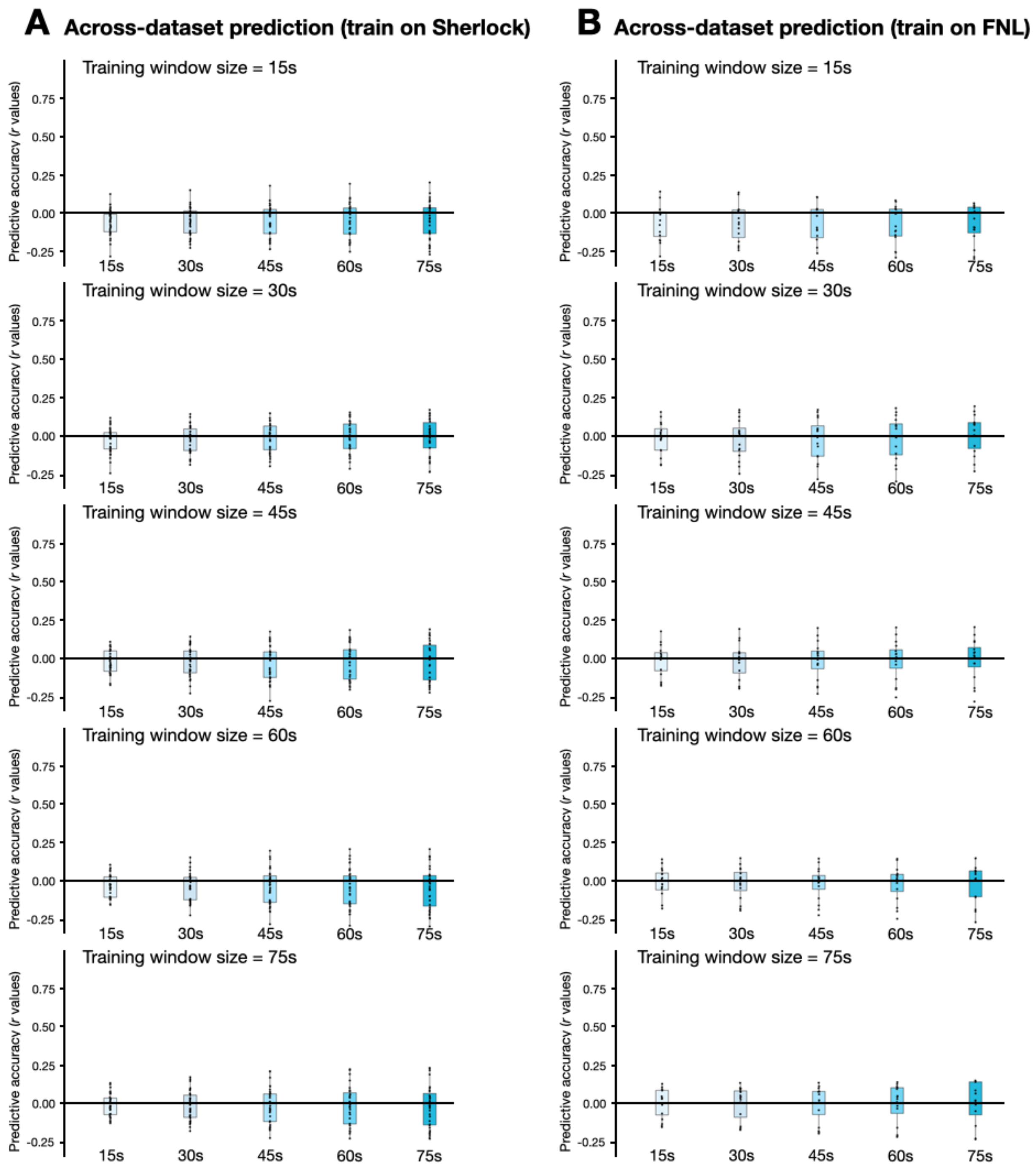


**Supplementary Figure 3. Across-dataset model predictive accuracy on valence as a function of sliding window size**, when training on *Sherlock* and testing on *Friday Night Lights* **(A)** and when training on *Friday Night Lights* and testing on *Sherlock* **(B)**. We extracted dynamic functional connectivity patterns from both *Sherlock* and *Friday Night Lights* using 5 different sizes of tapered sliding window: 15s, 30s, 45s (the original size in the main analysis), 60s and 75s, and tested each model on test data at each window size, resulting in 50 conditions (2 datasets x 5 window sizes at training x 5 window sizes at testing). None of the 50 conditions showed above chance predictive accuracy in predicting valence.

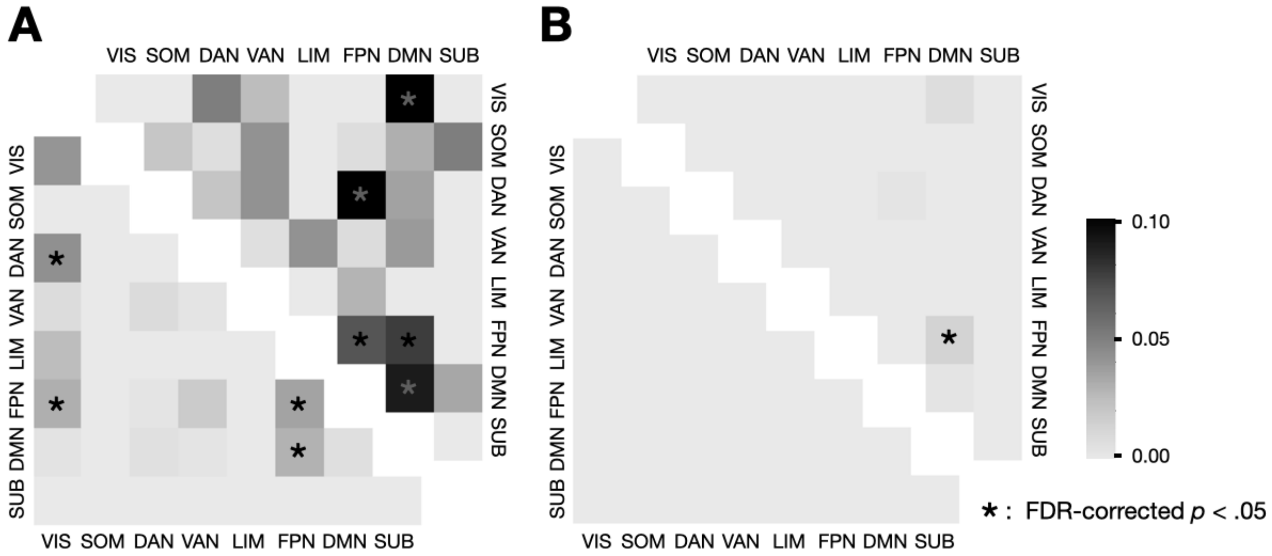


**Supplementary Figure 4. Functional anatomy of the overlap network between arousal and engagement.** Engagement network acquired from Song et al., 2021. In each figure, the upper triangle represents the network which contains functional connections that positively correlate with both arousal and engagement. The lower triangle represents the network which contains functional connections that negatively correlate with both arousal and engagement. Each cell represents the proportion of selected FCs among all possible FCs between each pair of functional networks. Networks pairs with above-chance selected FCs are indicated with an asterisk (one-tailed *t*-test, fdr-corrected $p < .05$). **A. The overlap between the arousal network and engagement network in the *Sherlock* dataset.** The connections between DMN and FPN, between DMN and VIS, between DAN and FPN as well as connections within the DMN and FPN positively predicted both arousal and engagement. The connections between VIS and DAN, between VIS and FPN, between FPN and DMN, as well as connections within the FPN negatively predicted both arousal and engagement (FDR-corrected $P < .05$). **B. The overlap between the *Sherlock-Friday Night Lights* arousal network and *Sherlock-Paranoia* engagement network.** Only the connection between DMN and FPN positively predicted both arousal and engagement in this cross-dataset manner.

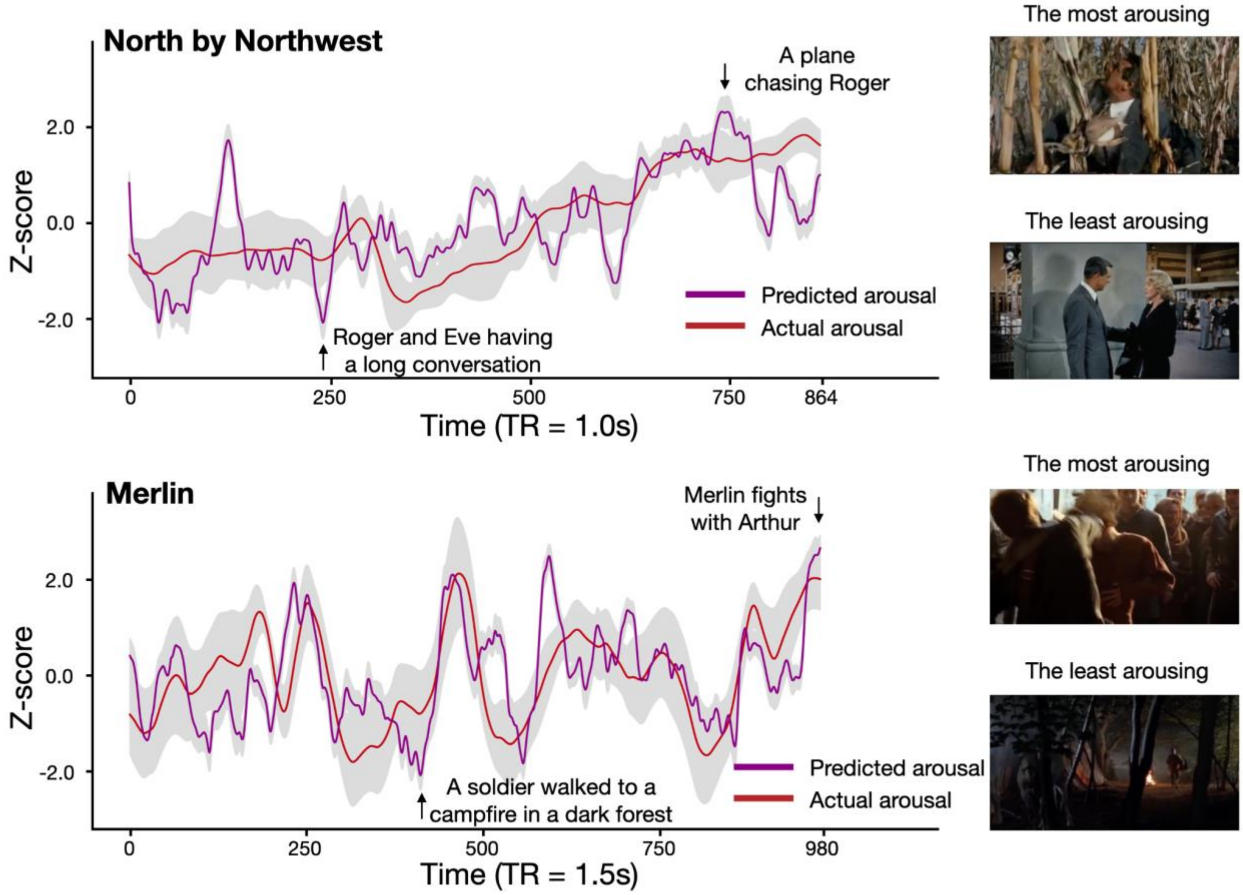


**Supplementary Figure 5. Arousal CPM predicts group-average arousal rating in two additional datasets.** Model predicted time courses across participants watching the same movie was averaged. This averaged predicted arousal time course significantly correlated with the actual group-average arousal rating. Model-predicted arousal time courses also corresponded with the plot of each movie. In *North by Northwest*, the most arousing moment predicted by the model occurred in the scene when the protagonist was being chased by a plane, while the least arousing moment occurred during a long conversation between characters; In *Merlin*, the most arousing moment predicted by the model occurred when the two main protagonists had a brawl in a tavern, while the least arousing moment occurred when a nameless soldier walked towards a campfire.

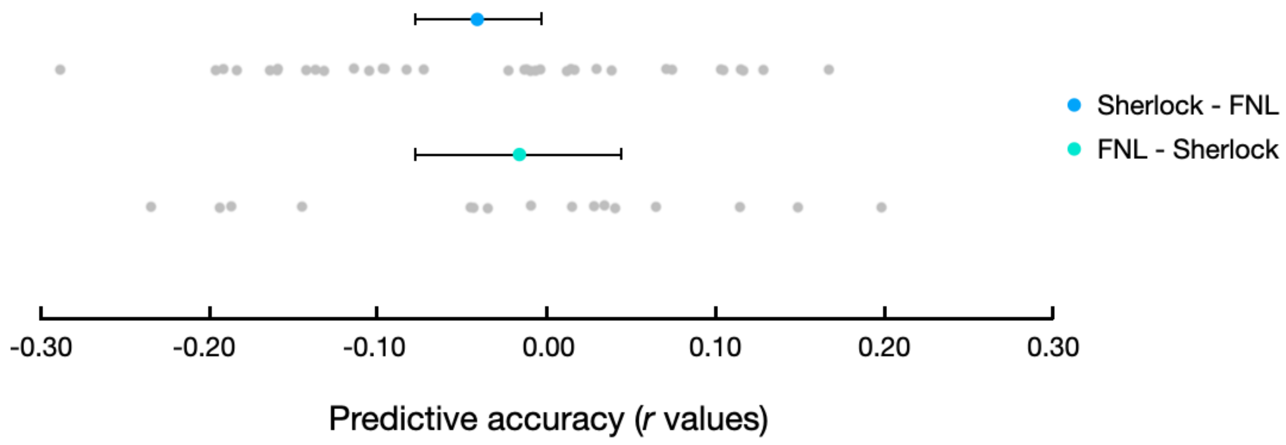


**Supplementary Figure 6.** Equivalence tests were performed to assess whether the across-dataset predictive accuracies in predicting valence fell within a predefined range around zero, which would suggest that they were not only statistically non-significant but also practically insignificant. We defined an equivalence interval of [-.100, .100], with the bounds determined based on a small effect size of r = .100. Each datapoint in the box plot represents the predictive accuracy of each round of cross-validation. The black horizontal lines show the 95% percent CI of the mean *r* value. The equivalence test was significant for both across dataset accuracies (*Sherlock* - *Friday Night Lights*: $p$ = .368, *Friday Night Lights* - *Sherlock*: $p$ = .306), indicating that the observed predictive accuracies were statistically indistinguishable from zero within the defined bound.

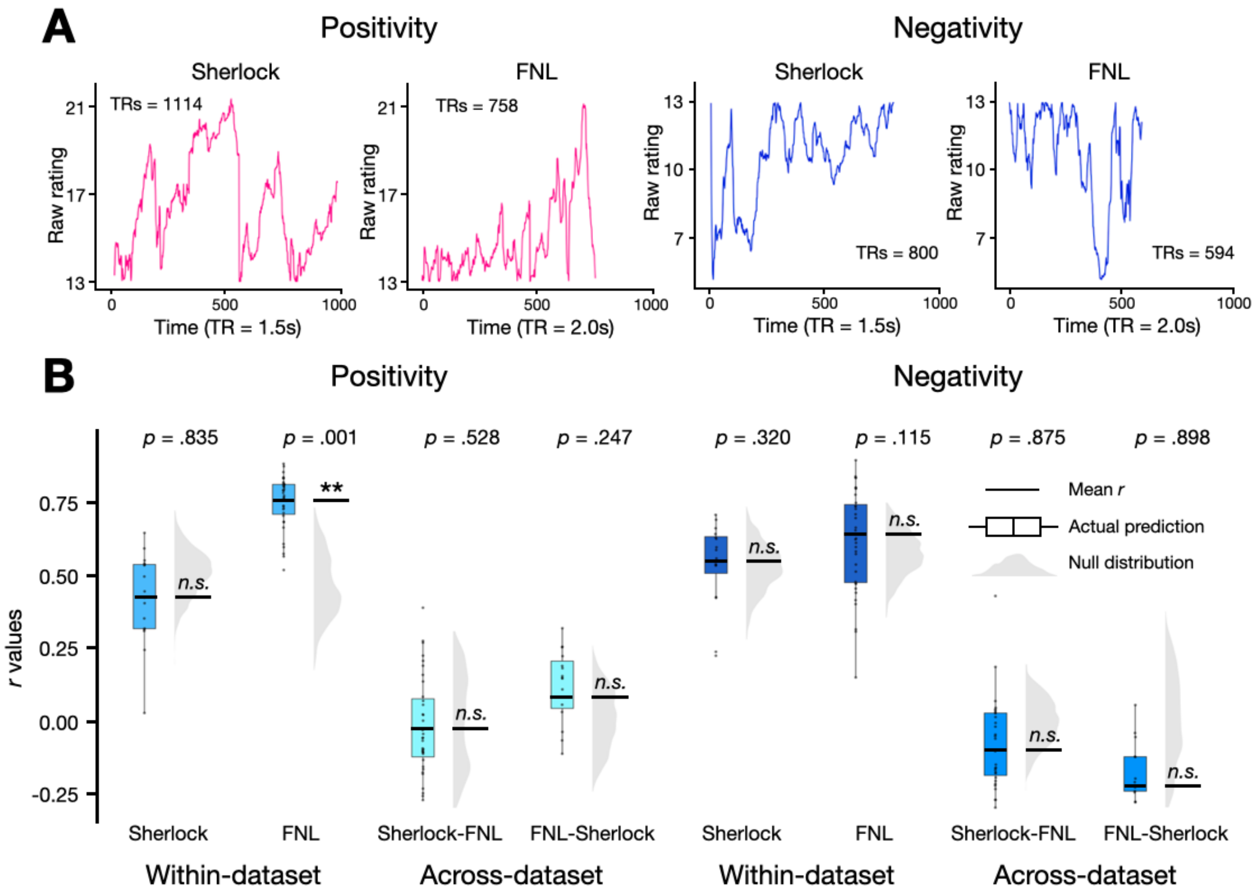


**Supplementary Figure 7. A.** Participants' subjective affective experience of positivity (left) and negativity (right) fluctuates over time during naturalistic movie watching. **B.** Dynamic functional connectivity does not predict subjective feelings of positivity and negativity. CPM performance in predicting positivity and valence for within-dataset (the left panel) and between-dataset (the right panel). The y-axis represents the predictive accuracy, as measured by Pearson's correlation between the model predicted time course and the observed group-average time course. Each datapoint in the box plot represents the predictive accuracy of each round of cross-validation. The black horizontal lines show the fisher-z transformed mean *r* value. The gray half-violin plots show the null distribution of 1000 permutations, generated by phase-randomizing the observed group-average before training and testing the models. *: $p < 0.05$, **:$p < 0.01$, n.s.: $p > 0.05$, as assessed by comparing the empirical mean predictive accuracy against the null-distribution.

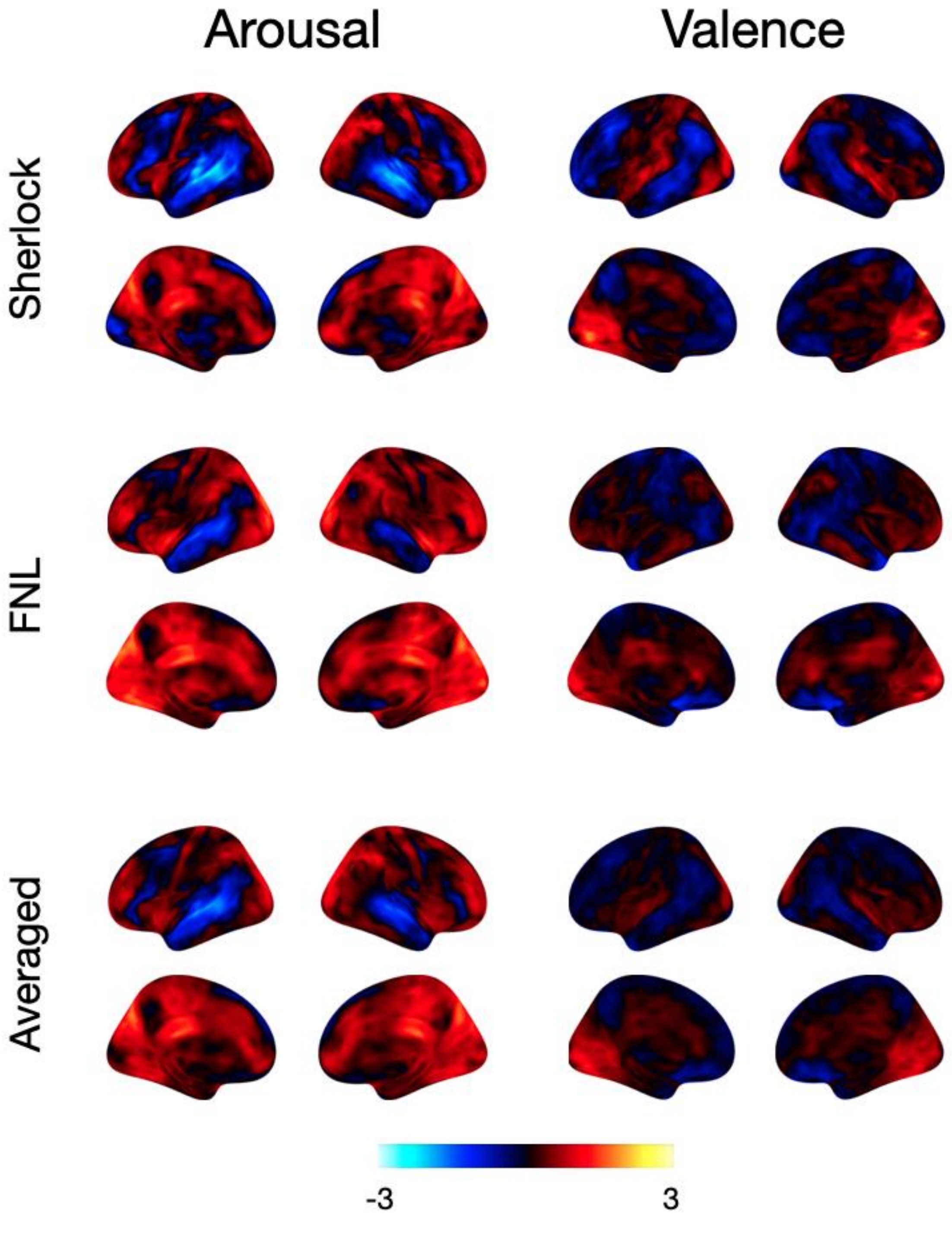


**Supplementary Figure 8. Unthresholded univariate parametric maps of arousal (left) and valence (right).** We built generalized linear models (GLMs) for *Friday Night Lights* and *Sherlock* separately to identify the voxels whose BOLD activity were associated with valence or arousal in each movie. The raw, unthresholded z-statistics are plotted here. Red indicates stronger activation during high-arousing moments, whereas blue indicates weaker activation during high-arousing moments.

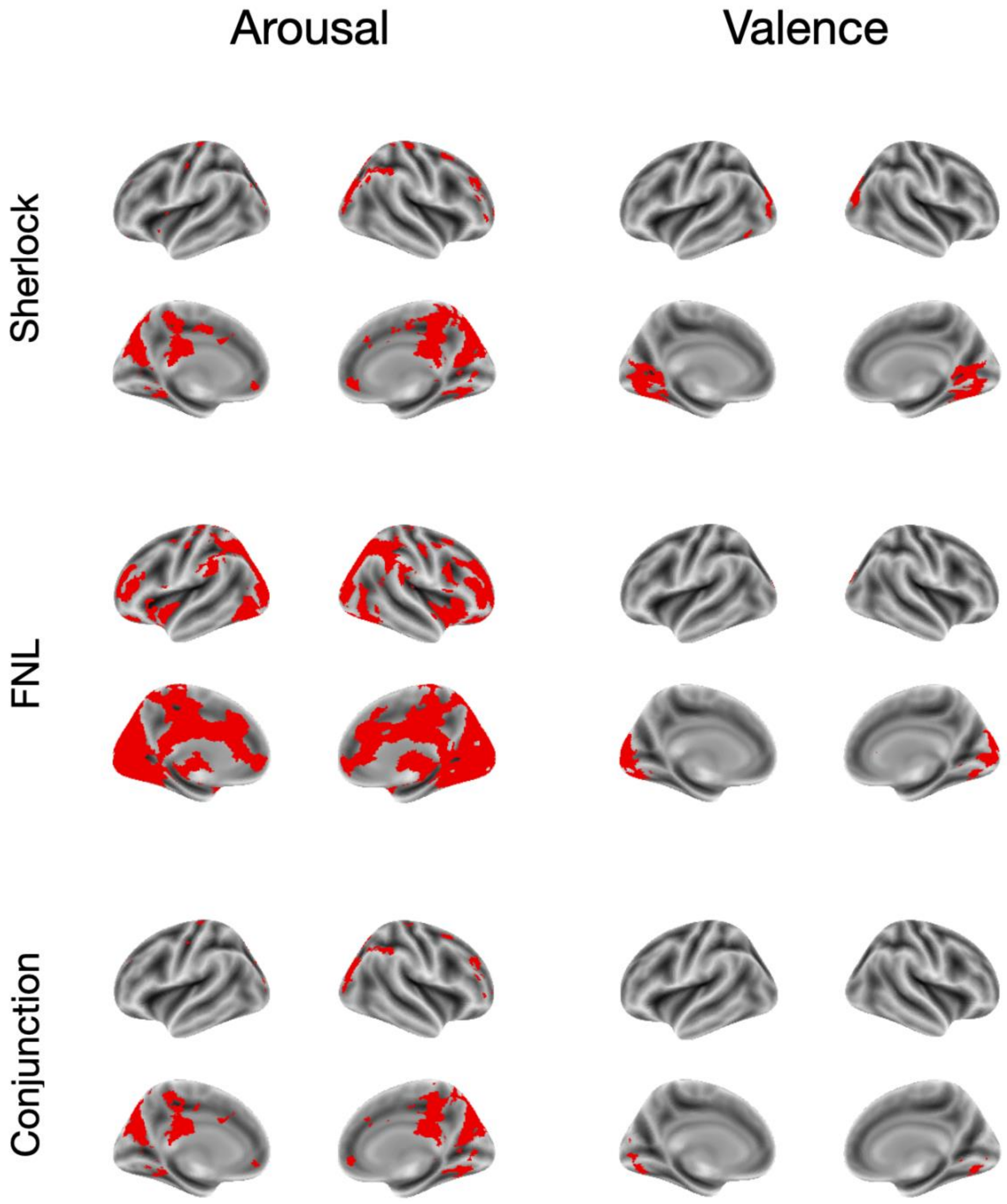


**Supplementary Figure 9. Thresholded univariate parametric maps of arousal (left) and valence (right) in the *Friday Night Lights* and *Sherlock* datasets, as well as the conjunction between the two datasets.** From the unthresholded z-statistics map, we performed a minimum statistic compared to the global null (MS/GN) test as the conjunction analysis to identify the voxels whose activations were consistently significantly related to valence or arousal across datasets (TFCE-corrected $p < .01$). We observed clusters of voxels that were more positively associated with arousal in the anterior cingulate gyrus, precuneus cortex, and insular. However, few voxels exhibit a significant positive association with valence.

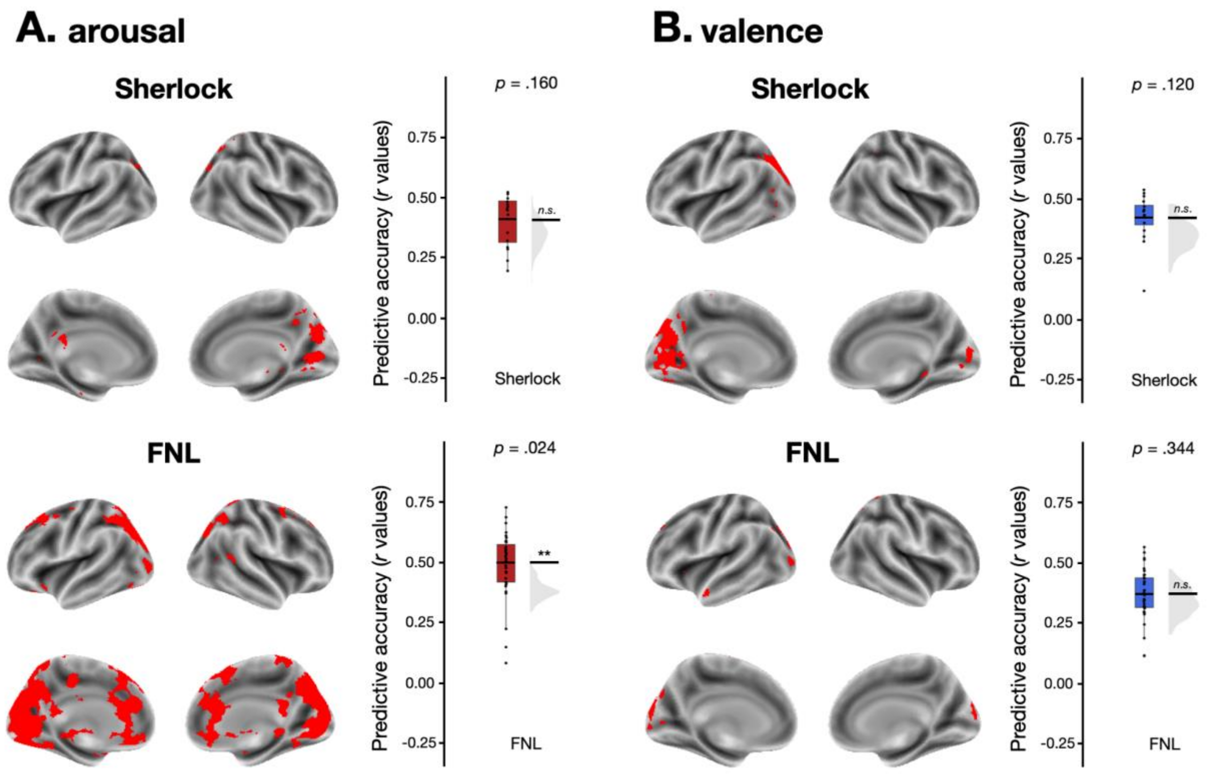


**Supplementary Figure 10. Multivariate patterns of brain activities do not predict the valence and arousal ratings within-dataset.** Multivariate-pattern based predictive model performance in predicting arousal **(A)** and valence **(B)** within-dataset. The brain map in each sub-plot indicates the voxels selected in the predictive modeling. The y-axis represents the predictive accuracy, as measured by Pearson's correlation between the model predicted time course and the observed group-average time course. Each datapoint in the box plot represents the predictive accuracy of each round of cross-validation. The black horizontal lines show the Fisher-*z* transformed mean *r* value. The gray half-violin plots show the null distribution of 100 permutations, generated by phase-randomizing the observed group-average before training and testing the models. *$p < 0.05$, **$p < 0.01$, n.s.: p > 0.05, as assessed by comparing the empirical mean predictive accuracy against the null distribution.
